# Reconstructing lineage-constrained gene-program dynamics with PhyloFM

**DOI:** 10.64898/2026.07.29.741462

**Authors:** Zihan Wang, Zhenyi Zhang, Xiaojing Yang, Chao Tang, Peijie Zhou

**Author notes:** Corresponding authors: (X.Y.), (C.T.), (P.Z.). These authors contributed equally (Zihan Wang, Zhenyi Zhang).

## Abstract

Cell-fate dynamics are defined not only by where cells end, but by when fate programs emerge, diverge, and reshape population structure. Much recent effort has been devoted to reconstructing these dynamics from temporally resolved single-cell RNA-seq and spatial transcriptomics. However, it is difficult because these measurements only provide destructive snapshots instead of continuous molecular histories. On the other hand, a diverse set of lineage-tracing methods are now available which could serve as temporal constraints to connect the dots. The challenge is how to utilize and unify these diverse ancestry records in the reconstruction of a continuous cell state dynamics. Here we introduce PhyloFM, a lineage-constrained flow-matching framework that maps heterogeneous lineage information onto a cell-fate-anchor topology and learns a continuous velocity field together with relative cell-abundance changes on a topology-regularized latent geometry. From the same integrated trajectories, PhyloFM predicts population transport, future fate probabilities, commitment timing and branch-resolved projected gene- and module-level dynamics. We tested PhyloFM across five lineage-tracing settings spanning *C. elegans* embryogenesis, LARRY hematopoiesis, pandaCREST ventral-midbrain development, zebrafish heart regeneration and spatial eTracer tumours. In *C. elegans*, our method improved population transport by 17.3% and lineage-grounded dynamic error by 16.3% relative to state-of-the-art dynamical baselines, with transport error reduced by 26.6% relative to additional lineage-aware control. Independent EPIC GFP reporter traces support the inferred timing of proneural, ciliated-neuron and late-selector programs, with 67% of matched reporter genes showing Pearson r > 0.6. On the spatial eTracer tumour benchmark, PhyloFM increased clone-grounded fate accuracy by 70.4% relative to the best spatial baseline. Together, these analyses show that diverse lineage records can serve as biological constraints for reconstructing continuous, interpretable molecular fate dynamics from single-cell snapshots.

## Introduction

During development, tissue repair, regeneration and disease progression, cell states evolve, commit to different fates, expand or decline and activate stage-specific molecular programs. Spatiotemporal single-cell atlases have provided increasingly quantitative data for these processes^1–4^ and have motivated recent efforts to build predictive virtual-cell models of how cell states change with time, context and perturbation^5–9^. Yet these measurements themselves remain discontinuous. Single-cell RNA-seq and spatial transcriptomic assays consume the cells they profile, so each sampled time point is a destructive snapshot of a different population. The continuous molecular histories that connect these snapshots—the routes cells take are not observed directly. This problem becomes even more serious when the cell state within a population bifurcates during the gap of these snapshots. Predictive models of cell states therefore require reconstruction of continuous dynamics beyond static-state catalogues.

Lineage tracing provides a natural source of biological prior information for this problem. By recording ancestry alongside molecular or spatial state, lineage experiments connect present cell states to observed clonal outcomes in embryogenesis, tissue repair, organoid differentiation and tumour evolution^10–16^. This additional modality can disambiguate cells that look similar transcriptionally but have different futures, and spatial lineage experiments further connect ancestry to local microenvironments^13,14,17–19^. However, the major obstacle is that lineage information is recorded in different forms across technologies. Barcode-based experiments can recover an inherited tag across sample time, allowing cells with the same barcode to be treated as members of the same clone over time. CRISPR-recording experiments often define relatedness among cells within one embryo, animal or organoid, although scars from independently sampled specimens typically lack a shared clone identity across time. Exact developmental trees provide fixed ancestry by position in a known tree, and clone-split or spatial lineage experiments add still different forms of future or tissue-context information. Therefore, these records are biologically related but technically difficult to compare within one unified analytical framework.

Existing computational methods address complementary parts of the continuous dynamics reconstruction problem^20,21^. Pseudotime^22–24^ and RNA velocity^25–34^ infer ordering or local molecular direction from expression profiles without using inherited future information to constrain population dynamics between destructive snapshots. For time-series datasets, optimal-transport^20,35–41^ and neural-dynamics approaches^42–55^ couple populations across sampled times and can learn continuous velocity fields and, in some cases, relative growth fields, but they usually rely on expression and time alone. Lineage-aware methods add a stronger biological prior. LineageOT^56^ and Moslin^57^ refine cross-time couplings, CoSpar^58^ infers transition maps and early fate bias, structural methods such as CARTA^10^, LinRace^59^, and LineageMap^60^ reconstruct differentiation maps, division histories or spatial lineage trees, and lineage-aware deep-learning or clone-embedding methods address clone-specific kinetics, fate prediction, historical reconstruction or clone-conditioned expression. While useful, they usually do not provide the same combination of a lineage-shaped representation, integrable velocity-growth dynamics and branch-resolved molecular-program dynamics in a unified manner. A general lineage-aware dynamical model should therefore translate diverse ancestry records into a common cell-state topology and use that topology to constrain the continuous paths, relative mass changes and molecular readouts inferred between snapshots.

Here we introduce PhyloFM, a lineage-topology flow-matching framework for reconstructing continuous molecular fate dynamics from destructive single-cell snapshots paired with diverse lineage-tracing records. PhyloFM separates topology construction from dynamical inference so that ancestry encoded in different experimental formats can constrain a shared dynamical model. Exact developmental trees are used directly, while clone, scar, clone-split and spatial lineage records are mapped by an appropriate topology constructor onto a shared graph of cell-state anchors built from comparable cell states across lineage-recording technologies. The graph also regularizes the latent geometry used for dynamical inference: straight latent bridges approximate short routes on the lineage-state topology and the intermediate states used by flow matching follow lineage-compatible paths. On this geometry, hybrid unbalanced optimal transport selects lineage-compatible couplings between adjacent snapshots, and simulation-free flow matching uses these coupled endpoints and topology-aligned bridges to learn continuous velocity and growth fields. When spatial coordinates are available, spatial regularization and spatial coupling costs extend the same geometry to local tissue structure. The anchor-graph reduction and the simulation-free flow matching objective build on lineage-state topology reduction^10^ and velocity-growth flow matching^43^, respectively. Thus, PhyloFM provides the topology-regularized latent geometry and hybrid lineage-aware coupling that place ancestry inside the representation on which continuous dynamics are learned. From the same integrated trajectories, PhyloFM predicts population transport, per-cell fate probabilities, commitment timing, relative cell-abundance change and branch-resolved gene- or module-level program dynamics. These molecular readouts are evaluated against observed descendants, curated regulatory programs and, where available, independent reporter measurements, so that lineage constrains the full dynamical path instead of serving only as a post hoc trajectory label.

We applied PhyloFM across five lineage-record regimes chosen to span the main ways ancestry is measured in current single-cell studies: the exact *C. elegans* embryonic lineage, shared-barcode hematopoiesis with LARRY, clone-split CRISPR barcoding in pandaCREST mouse ventral-midbrain development, per-animal CRISPR scars in zebrafish heart regeneration and spatial clone families in eTracer tumours. Across these settings, PhyloFM improves population transport and clone-grounded fate prediction relative to dynamical baselines, with additional lineage-aware controls in the exact-lineage benchmark. The *C. elegans* analysis provides the strongest molecular validation: inferred proneural, ciliated-neuron and late-selector programs agree with independent EPIC GFP reporter dynamics that were not used for training, supporting the ability of lineage constraints to recover biologically timed molecular programs from destructive snapshots. The remaining datasets show that the same framework recovers clone-grounded hidden-future bias, scar-linked repair branches and spatially organized tumour-state progression across distinct lineage-record formats. Together, these results show that heterogeneous lineage records can serve as experimental constraints for reconstructing continuous, interpretable and testable molecular fate dynamics.

## Results

### Overview of PhyloFM

PhyloFM formalizes lineage information as a technology-agnostic topology over cell-state anchors. Exact tree edges, shared barcodes, clone-split families, CRISPR scars and spatial clone-family labels are each converted into this anchor graph, which preserves the ancestry relationships that are comparable within the corresponding experiment. In the experiments below, exact trees supply the graph when available. For non-exact records, CARTA-derived summaries provide one implementation of the required anchor graph, and the PhyloFM objective acts on the resulting graph distances. The graph therefore becomes a common coordinate system linking lineage technologies to the dynamical model (Fig. 1a). During representation learning, cells linked by short graph paths are placed nearby in latent space, and cells separated by long paths are separated up to a scaled margin. This topology regularization turns the latent space into a geometry of admissible lineage transitions. Endpoint coupling determines which observed populations should be connected across sampled time points. The topology-shaped latent space determines the route taken between those endpoints, so straight interpolation in latent coordinates approximates a short path on the lineage-state graph and the states sampled for flow matching inherit the same lineage constraint.

**Figure 1:**
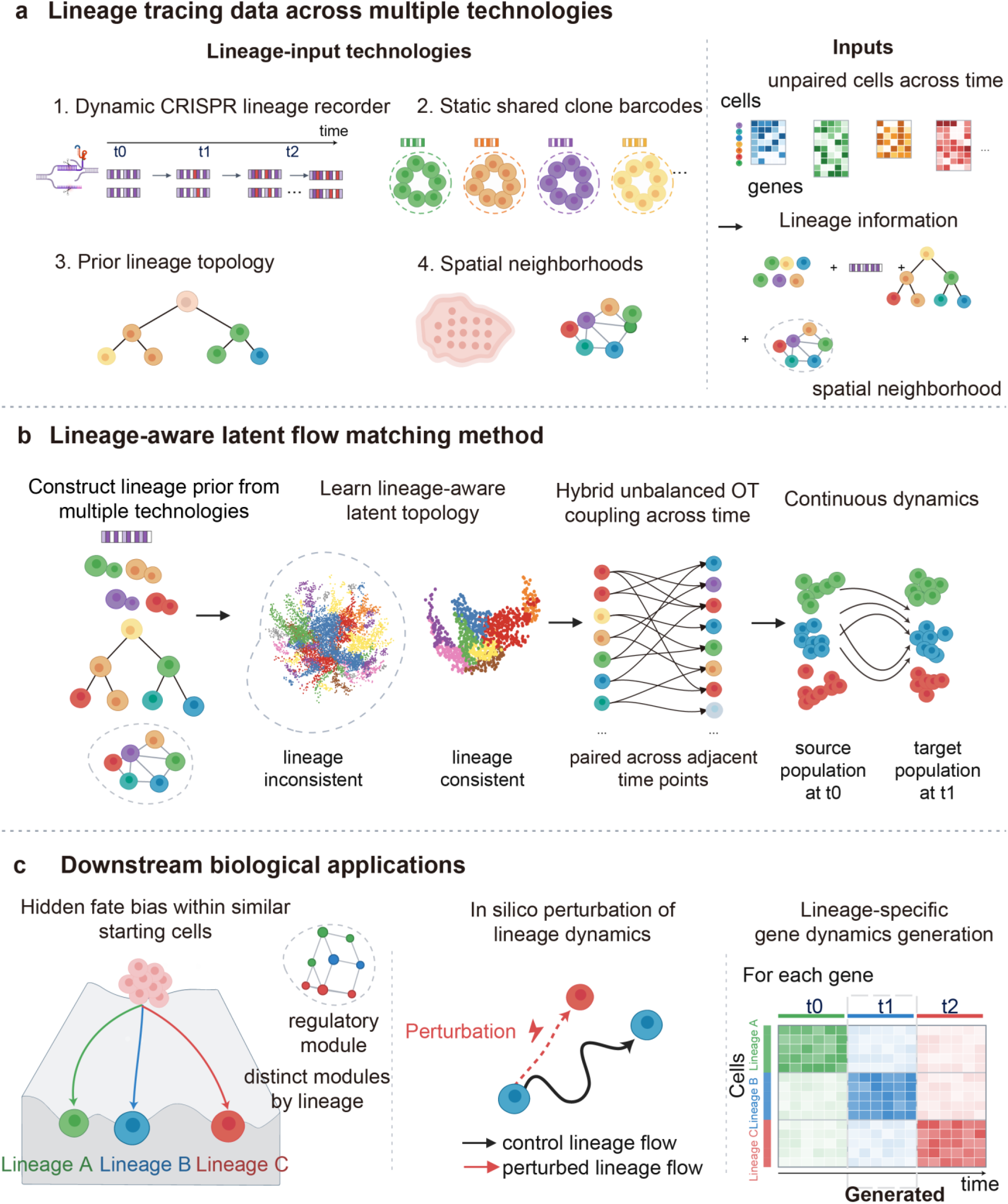
PhyloFM integrates heterogeneous lineage records into continuous cell-state dynamics. a, Lineage-tracing studies combine temporally resolved single-cell transcriptomes, and optionally spatial coordinates, with ancestry information generated by dynamic CRISPR recorders, shared clone barcodes, prior lineage topologies or spatial neighbourhood graphs. These inputs provide gene-expression matrices across time together with lineage and spatial constraints. b, PhyloFM first converts each technology-specific record into a lineage prior, then learns a lineage-aware latent topology, computes hybrid unbalanced optimal-transport couplings between adjacent time points, and trains simulation-free velocity and growth fields over the latent space. c, The resulting model supports hidden-future inference among similar starting cells, in silico perturbation of lineage dynamics and reconstruction of lineage-specific gene programs.

Continuous dynamics are learned on this lineage-shaped geometry. Adjacent time points are coupled by a hybrid optimal-transport problem whose cost measures whether two cells are plausible descendants in molecular state, lineage-state position and, when available, tissue location. Because the coupling is unbalanced, its row mass provides the per-cell growth target used to train a growth network alongside the velocity field. Flow matching then converts the lineage-aware paired endpoints into a continuous vector field on topology-aligned latent bridges. The resulting field predicts unsampled intermediate states while training on points that remain aligned with the lineage-state graph. At inference, latent state and cell weight are integrated jointly, and the latent trajectory is decoded to gene-expression profiles. Population transport, per-cell fate probabilities, growth estimates, gene-dynamics trajectories and module-perturbation responses are then derived from the same learned trajectory. This yields a coupled velocity-growth model in which endpoint matching, intermediate trajectories, population expansion and molecular programs are all constrained by the lineage topology (Fig. 1b).

We evaluated PhyloFM on five lineage-tracing datasets that together cover major forms of lineage recording currently in use. The exact embryonic lineage of *C. elegans* provides a stringent benchmark because its known tree supports evaluation of population transport, lineage-local dynamics, route structure and molecular programs^61,62^. The LARRY hematopoiesis dataset gives a clone-grounded fate benchmark through shared lentiviral barcodes that label founder cells with their later myeloid outcomes^13^. pandaCREST links E11.5 progenitors to day-7 organoid derivatives by clonal splitting^17^, exposing hidden future within a coarse progenitor state. These two datasets test whether reusable clone information can be converted into predictive fate dynamics before terminal states are evident. The zebrafish and eTracer datasets test portability beyond reusable clone labels. Zebrafish heart regeneration is a multi-animal repair series in which CRISPR scars are generated separately in each fish^63^, so cross-time matching must pass through shared cell-state anchors. eTracer then extends the evaluation to an emerging spatial-lineage setting in which tumour clones, tissue location and transcriptome are measured together^19^. Across this hierarchy, we report quantitative comparisons, lineage ablations and biological analyses that ask whether ancestry can make hidden future states visible before they are obvious in expression space (Fig. 1c).

Because cell state, relative mass-balance and molecular-program readouts are derived from the same integrated trajectories, PhyloFM makes a directly testable prediction: inferred molecular dynamics at unobserved or intermediate states should agree with independent measurements in timing and branch specificity. We test this most directly in *C. elegans*, where an exact embryonic lineage provides transition-level ground truth and independent EPIC fluorescent-reporter time series provide orthogonal reporter-based validation of gene-program timing, before applying the same framework across four additional lineage-record regimes.

### Exact-lineage and reporter benchmarks support branch-resolved molecular dynamics in *C. elegans*

*C. elegans* provides a stringent setting for asking whether lineage-constrained dynamics can recover molecular fate programs from destructive snapshots. Its invariant embryonic lineage places every sampled cell on a known developmental tree^56,57,61,62^, providing exact ancestry for local transition benchmarks, while the independent EPIC fluorescent-reporter atlas provides regulator time series that were not used for training. We trained PhyloFM on expression and the exact lineage record of the ABpxp founder pool across seven time points (170, 210, 270, 330, 390, 450 and 510 min), and used this system to ask three questions: whether lineage-shaped dynamics improve population transport and exact-lineage local transitions; whether accurate lineage matching alone is sufficient, or whether the learned geometry adds information beyond endpoint matching; and whether inferred molecular trajectories agree with independent reporter measurements, providing validation beyond lineage-consistent transport.

Raw expression did not organize ABpxp cells by developmental order. Cells from different lineage depths and sub-branches were intermingled in the UMAP, so the developmental sequence encoded by the tree was only weakly visible in the expression manifold (Fig. 2a). In PhyloFM’s topology-regularized latent space, the same cells formed a more ordered geometry in which nearby positions followed ABpxp progression and continuous trajectories traced lineage-shaped routes from founders toward descendants (Fig. 2b). The geometry translated into quantitative gains against four lineage-blind dynamical baselines spanning the main design choices in this field: OT-CFM^64^, VGFM^43^, scDiffEq^54^ and TIGON^46^. For non-holdout transport, PhyloFM reduced W1 by 17.3% and W2 by 12.3% relative to the best non-PhyloFM baseline, reaching W1 = 5.11 and W2 = 5.54 compared with the best baseline values of 6.18 and 6.32. In the time-point hold-out test, where each intermediate stage was omitted in turn, PhyloFM also gave the lowest W1 and W2 (6.24 and 6.46). Lineage-local dynamic error, which scores model predictions against tree-connected descendants, was also lowest for PhyloFM. It reached 41.97, a 16.3% reduction relative to VGFM at 50.13, followed by scDiffEq at 59.98, OT-CFM at 67.78 and TIGON at 68.40 (Fig. 2c). Removing the topology, shuffling lineage labels or rebalancing the topology–reconstruction terms each shifted the model into a higher-error regime, confirming that these gains depend on the lineage signal itself (Supplementary Fig. S1f).

**Figure 2:**
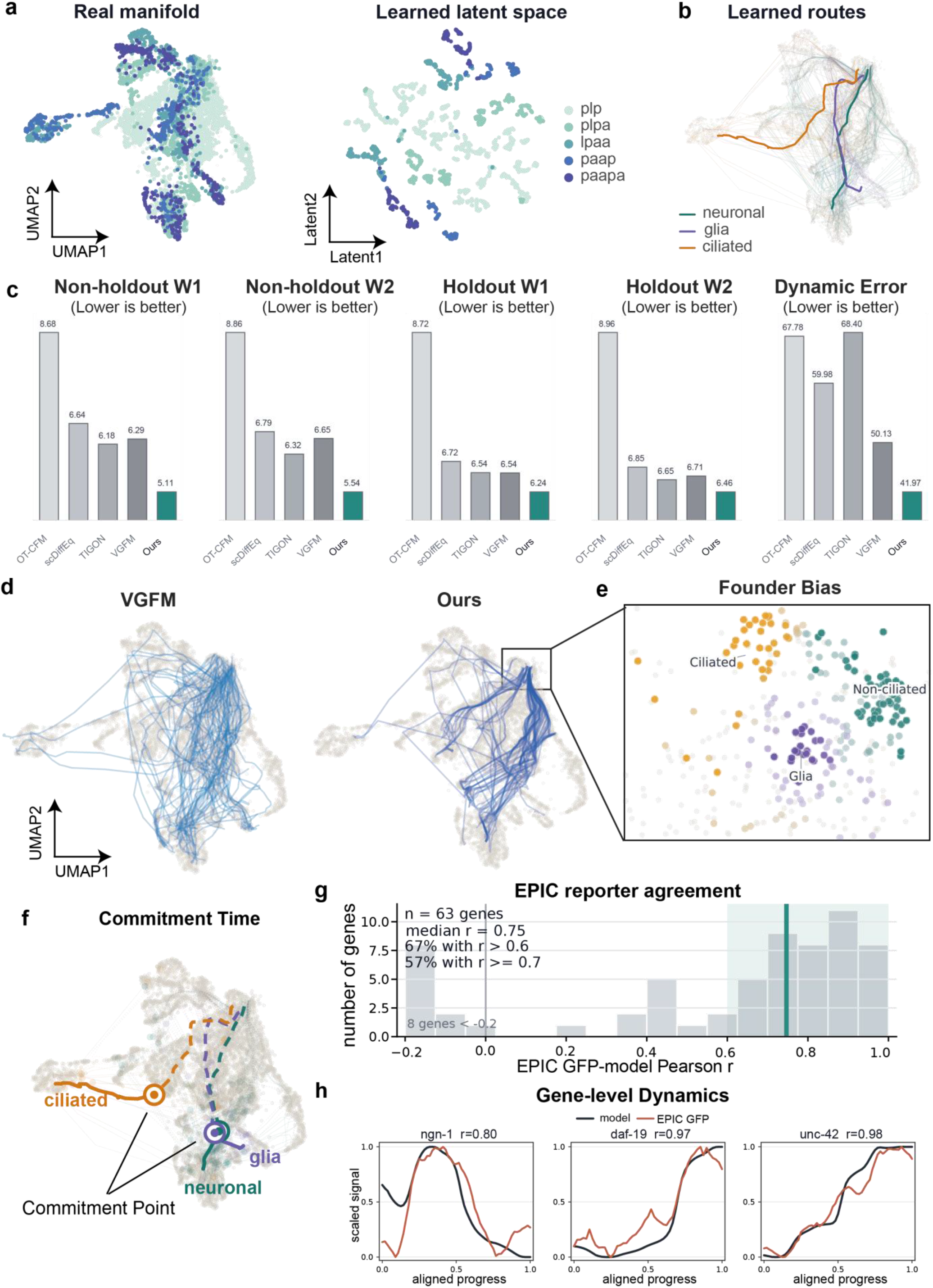
Exact-lineage benchmarking and route programs in *C. elegans*. a, Real expression manifold and PhyloFM latent representation of ABpxp-lineage cells, coloured by retained exact-lineage prefixes. b, Learned routes from the ABpxp founder pool toward neuronal, glial and ciliated futures. Thin curves show individual inferred trajectories and thick curves show route-level summaries. c, Quantitative comparison with baseline dynamical models using non-holdout W1, non-holdout W2, time-point holdout W1, time-point holdout W2 and lineage-local dynamic error; lower values indicate better agreement with observed distributions or lineage-defined local transitions. d, Route geometry comparison between VGFM and PhyloFM in the shared founder-region view. e, Founder-bias map identifying early ABpxp cells biased toward ciliated, glial or non-ciliated neuronal futures before terminal identities are reached. f, Route-specific commitment-time map, with marked commitment points indicating when each future becomes dominant. g, Distribution of EPIC GFP reporter-model Pearson correlations across genes with usable EPIC2 reporter traces and inferred gene trajectories. h, Inferred dynamics of *ngn-1*, *daf-19* and *unc-42* compared with matched EPIC GFP reporter profiles measured in live *C. elegans* embryos. Black curves show PhyloFM predictions, red curves show EPIC GFP signals, and titles report Pearson correlations.

We next isolated the contribution of lineage-aware endpoint matching. Because the four dynamical baselines did not use lineage information during training, we evaluated Moslin-FM, a lineage-aware control that used Moslin-derived matchings as training pairs for the same flow-matching model without representation learning. The static coupling used by PhyloFM had slightly lower static-coupling error than Moslin, LineageOT and control couplings (normalized error 0.33 versus 0.36, 0.46 and higher; Supplementary Fig. S2a). More importantly, Moslin-FM approached PhyloFM in lineage-local dynamic error (42.9 versus 42.0) but retained substantially higher distributional transport error (W1 = 7.0 and W2 = 7.5 versus 5.1 and 5.5; Supplementary Fig. S2b). Thus, while lineage-aware endpoint matching captured local lineage transitions, PhyloFM further transformed the static coupling into valid continuous dynamics. Time-resolved curves confirmed that this advantage was sustained across the full developmental series (Supplementary Fig. S1a-e). Ablation analysis showed that removing lineage information, shuffling lineage labels, or altering the topology-reconstruction balance each shifted the model into a higher-error regime (Supplementary Fig. S1f).

We next asked whether these state-space gains carried decodable molecular information. We compared inferred gene trajectories with EPIC GFP reporter time series that were not used for training^65^, topology construction or model selection. With bounded-lag alignment to accommodate reporter timing differences, inferred trajectories showed overall agreement with the reporter profiles across 63 genes with usable EPIC traces (median Pearson r = 0.75; 67% of matched genes with r > 0.6; Fig. 2g). The anchor regulators were individually well supported: ngn-1, daf-19 and unc-42 reached reporter–model correlations of 0.80, 0.97 and 0.98, respectively (Fig. 2h). Additional high-agreement examples included unc-86 (r = 0.99) and several selector or developmental regulators with correlations between 0.91 and 0.94 (Supplementary Fig. S6). This independent reporter agreement is consistent with the inferred trajectories capturing biologically timed molecular programs in addition to lineage-consistent transport.

With transport, local dynamics and reporter-supported molecular timing established, we used the same trajectories to resolve route-specific fate programs. Within the ABpxp founder pool, PhyloFM separated cells into future ciliated-, neuronal- and glia/excretory-biased domains before terminal identities were evident, whereas the lineage-blind VGFM baseline produced more diffuse routes over the same manifold (Fig. 2d,e and Supplementary Fig. S3). With commitment defined as the point at which the future route became dominant, ciliated sensory-neuron routes consolidated earliest, other neuronal routes followed, and glial routes sharpened most gradually (Fig. 2f). This ordering is consistent with established features of *C. elegans* development, including early differentiation and ciliogenesis of ciliated sensory neurons, together with later neuronal maturation and dendrite-development programs^66–69^. Grouping decoded genes by temporal profile along the inferred routes recovered three programs matching this commitment order. A cilia-glia buffer pattern, exemplified by daf-19, rose specifically along the ciliated route and was suppressed on the neuronal route. A shared neurogenic-priming pattern, exemplified by ngn-1 (alongside ztf-11), peaked early on the ciliated and neuronal routes and then decayed. A late-selector pattern, exemplified by unc-42 (alongside dma-1 and fmi-1), turned on last and was strongest on the neuronal terminal (Supplementary Fig. S4). This is consistent with established *C. elegans* biology, in which daf-19 controls ciliogenesis in sensory neurons^70^, ngn-1 acts as an early proneural factor^71^, and unc-42 functions as a terminal neuronal selector^72^. We then compared the generated branches with a set of *C. elegans* nervous-system regulatory relationships described in Supplementary Note S3.4. PhyloFM recovered 70% of the documented regulator-to-program directions, compared with 53% for the best baseline, with examples spanning GABAergic identity, ciliogenesis, cholinergic motor identity, touch-receptor specification and late neuronal maturation (Supplementary Fig. S5). We interpret these directionality analyses as program-level support and hypothesis generation, whereas the exact-lineage local dynamics and EPIC reporter comparisons provide the direct quantitative validation of molecular timing.

Together, the *C. elegans* benchmark shows that lineage-constrained dynamics improve population transport, preserve exact-lineage local transitions beyond what lineage matching alone achieves and reproduce independently measured reporter timing. This establishes PhyloFM’s central capability before testing its portability across lineage-record technologies.

### Clone-grounded fate prediction in hematopoietic founders

The LARRY hematopoiesis dataset provides a complementary benchmark because lentiviral barcodes are inherited during hematopoietic differentiation and can be read across sampled days, so cells sharing a barcode provide direct clone-level evidence for future fate (Fig. 3a)^13^. This design allows fate prediction to be evaluated against inherited clone outcomes and tests whether the learned geometry exposes clone-defined future bias before differentiation. We trained PhyloFM on day-2 founders and later differentiated states, using the LARRY barcodes to evaluate future monocyte and neutrophil bias. The same system also gives a biological check on the learned dynamics, because monocyte-neutrophil fate choice is linked to well-studied myeloid regulators. The global hematopoietic landscape organized cells from an undifferentiated region toward monocyte, neutrophil, basophil, mast, megakaryocyte, erythroid and lymphoid states, with PhyloFM-generated trajectories tracing continuous paths over this manifold (Fig. 3b).

**Figure 3:**
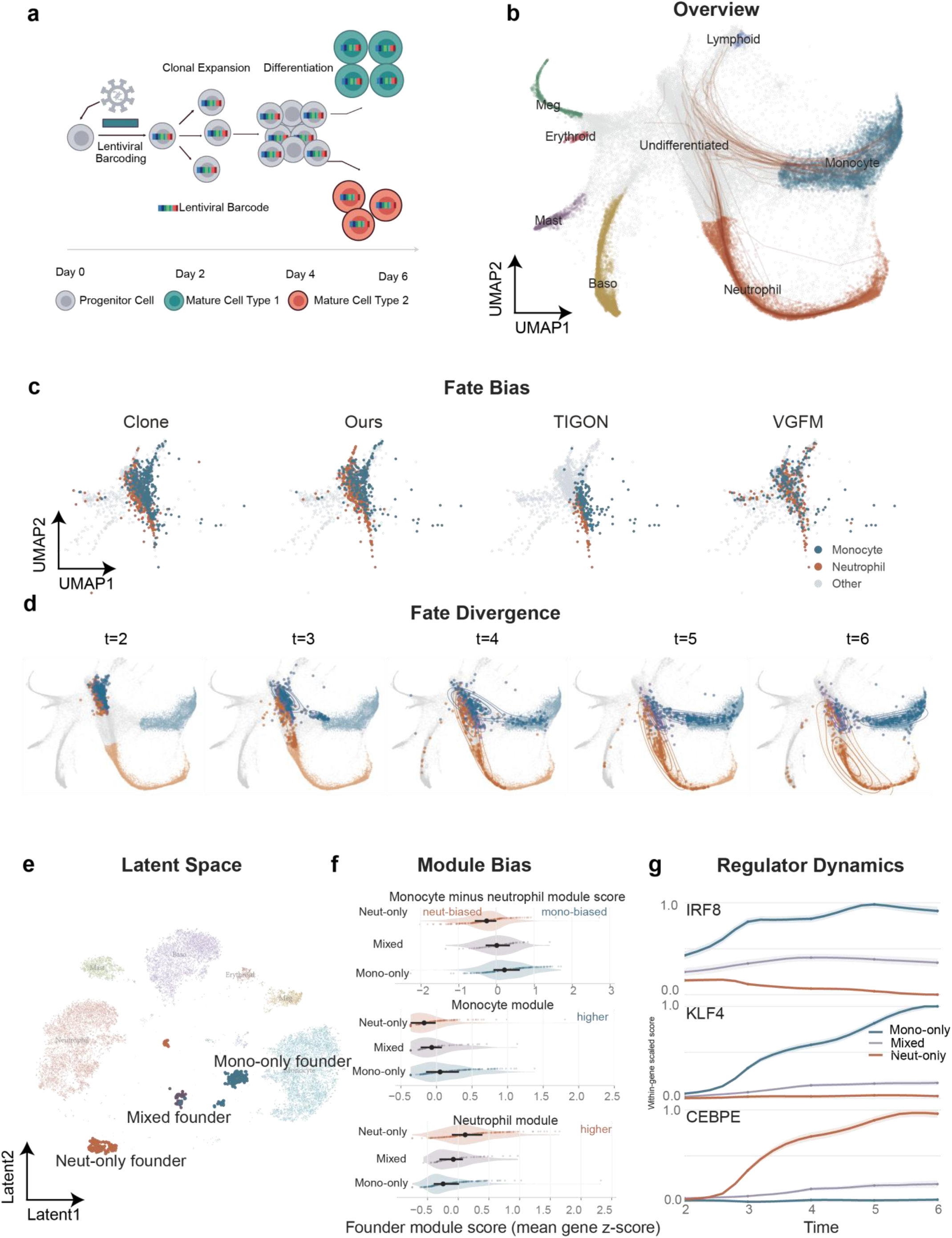
Clone-grounded fate prediction in LARRY hematopoiesis. a, Lentiviral barcoding records clonal expansion and differentiation from hematopoietic progenitors to mature blood lineages across the LARRY time course. b, Global cell-state manifold with PhyloFM trajectories from undifferentiated cells toward differentiated myeloid lineages. c, Founder fate-bias maps comparing clone-derived labels, PhyloFM predictions and baseline predictions for monocyte- and neutrophil-biased founders. d, Generated trajectories show progressive divergence of monocyte- and neutrophil-biased founder populations over time. e, PhyloFM latent space separates mono-only, mixed and neut-only founder regimes. f, Founder module scores for monocyte-minus-neutrophil, monocyte and neutrophil programs across clone-defined regimes. g, Regulator dynamics for *IRF8*, *KLF4* and *CEBPE* along mono-only, mixed and neut-only trajectories.

Barcode-derived labels separated day-2 founders into monocyte-biased, neutrophil-biased and other cells. PhyloFM recovered the arrangement of the monocyte and neutrophil founder domains more closely than other baselines and achieved the highest LARRY fate accuracy (Fig. 3c and Supplementary Fig. S7), indicating that the model recovers meaningful founder bias from clone information. The model also inferred where population expansion was concentrated along the hematopoietic landscape. Predicted growth was highest in progenitor-dominated earlier time points and lower later (Supplementary Fig. S8a). In silico perturbation of individual genes highlighted inflammation- and cytokine-associated regulators including *Nfkbia*, *Irf8* and *Il6st* that reduced the predicted expansion signal, while phagolysosome- and granule-related genes such as *Ctss*, *Cd63* and *Srgn* increased it (Supplementary Fig. S8b). Genes associated with this growth signal were enriched for inflammatory-response, cytokine-mediated signalling and mononuclear-cell migration terms (Supplementary Fig. S8c), placing the learned growth field on a known inflammation-hematopoiesis interface^73,74^.

Generated trajectories showed how this hidden founder bias resolves over time. Monocyte-biased and neutrophil-biased trajectories diverged from a shared progenitor region from day 2 toward day 6 (Fig. 3d). These clone-defined founder groups were largely intermingled in the expression manifold, but separated clearly in PhyloFM’s learned latent space, with mono-only, mixed and neut-only regimes occupying distinct regions (Fig. 3e). To ask whether these founder regimes differ at the molecular level, we scored each founder on monocyte and neutrophil modules assembled from canonical lineage regulators and differentiation markers. Mono-only founders were shifted toward the monocyte module, neut-only founders toward the neutrophil module, and mixed founders occupied an intermediate range (Fig. 3f and Supplementary Fig. S9a), indicating that clone-defined future bias was already visible in the founder population^75,76^. Regulator dynamics along generated trajectories mirrored this separation, with *Irf8* and *Klf4* rising along mono-only trajectories and *Cebpe* rising along neut-only trajectories (Fig. 3g). Gene-correlation and enrichment analyses linked the monocyte side to inflammatory and interferon-related programs and the neutrophil side to secretory-granule organization and myeloid differentiation (Supplementary Fig. S9b,c), consistent with established roles for *IRF8*, *KLF4* and *CEBPE* in myeloid fate control^77–79^. To test this molecular structure at a larger scale, we compared generated trajectories with a literature- and database-grounded myeloid regulatory benchmark described in Supplementary Note S3.4. PhyloFM recovered 81% of expected regulator-to-program directions, exceeding TIGON at 74% and all other comparators, with representative relationships involving *Irf8*, *Klf4* and *Cebpe* across monocyte and neutrophil programs (Supplementary Fig. S10). In silico perturbation of the monocyte and neutrophil programs in founder cells shifted terminal fate fractions in the expected directions (Supplementary Fig. S9d), in line with the regulator-to-fate associations^54,80^. Together, the analysis on the hematopoietic dataset shows that PhyloFM uses barcode-certified clone outcomes to resolve early monocyte-neutrophil bias and connects this bias to a directional myeloid regulatory program before terminal fate is fully established.

### Hidden fate structure linked to branch-specific regulatory programs

pandaCREST (progenitor and derivative associating CREST) is a CRISPR-based lineage-tracing strategy in which Cre-dependent Cas9 activation drives stochastic editing of two genomic recorder arrays (V1 and V2), generating highly diverse and stably inherited barcodes in a defined cell population^17^. In this study, barcoding was targeted to En1-Cre-labelled E11.5 mouse ventral-midbrain progenitors. Each clone was then split, with one part profiled in vivo at E11.5 and the other grown for one week as a brain organoid and profiled at day 7. Shared V1+V2 barcodes therefore connect source progenitors to organoid derivatives, making pandaCREST a direct test of whether clone-split ancestry can expose future bias within a progenitor state that is still transcriptionally coarse (Fig. 4a,b).

**Figure 4:**
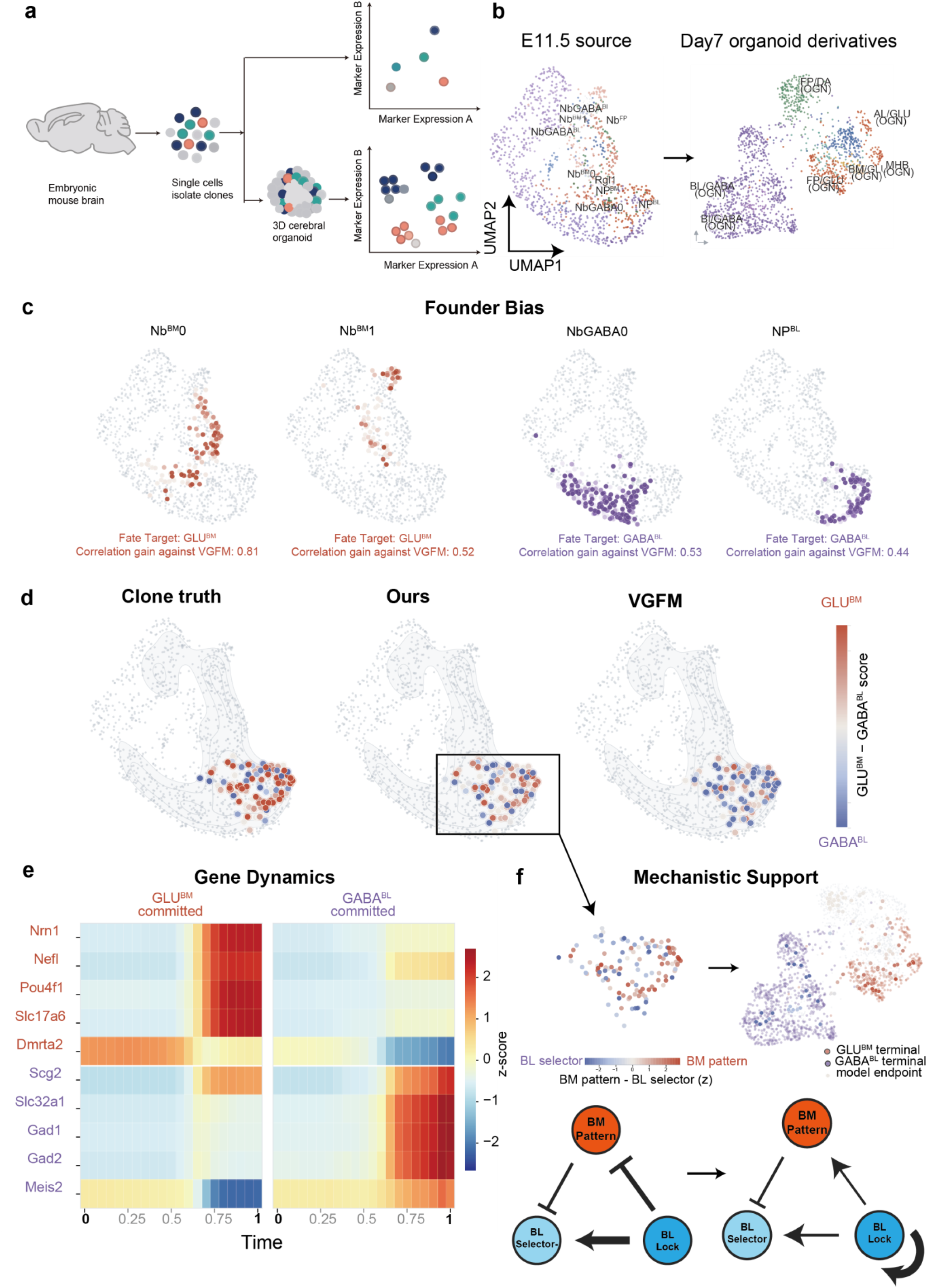
Clone-split pandaCREST reveals hidden branch bias in embryonic brain progenitors. a, pandaCREST links E11.5 embryonic mouse-brain source cells to day-7 organoid derivatives through clone splitting and inherited CRISPR barcodes. b, Source and derivative manifolds showing E11.5 progenitor states and organoid outcomes. c, Founder-bias maps identify source regions enriched for BM/GLU- or BL/GABA-directed futures, with correlation gains over VGFM shown for representative progenitor states. d, Same-state hidden-future benchmark for NPBM progenitors, comparing clone-derived fate margins with PhyloFM and VGFM predictions. e, Gene-expression dynamics along BM/GLU-committed and BL/GABA-committed trajectories. f, Mechanistic support for the inferred branch-control program, including BM-pattern, BL-selector and BL-lock modules and their inferred regulatory interactions.

At the coarse level, clone labels recovered the expected source-to-derivative structure. NPBL sources aligned with GABA^BL^-like derivatives, NPBM sources aligned with GLU^BM^-like derivatives, and Rgl1 aligned with the DA^FP^ side. Within the E11.5 source manifold, PhyloFM identified founder regions with strong future enrichment, including NbBM0 and NbBM1 toward GLU^BM^ outcomes and NbGABA0 and NPBL toward GABA^BL^ outcomes (Fig. 4c), in line with the principal lineage axis in the original study.

The more stringent test was the NPBM compartment, where cells share a coarse source annotation but their clones split between GLU^BM^-biased and GABA^BL^-biased outcomes. Across 127 NPBM cells, the fate bias predicted by PhyloFM tracked the clone-observed bias closely (Spearman ρ = 0.68, Pearson r = 0.71), while VGFM showed weak agreement with this same-state future bias (Fig. 4d). Thus, the lineage-shaped representation recovered a future axis that was present in the clone record before clear separation in expression space.

We next asked what molecular programs define this clone-aligned axis. Genes most associated with the GLU^BM^ or GABA^BL^ future marked different regions of the NPBM founder manifold, showing that the two future-biased groups already carried distinct molecular tendencies at the source stage (Supplementary Fig. S11a-d). Enrichment analysis linked the GLU^BM^ side to reduced motility and migration terms, and linked the GABA^BL^ side to transcriptional regulation and nervous-system development terms (Supplementary Fig. S11e). Along generated trajectories, GLU^BM^-committed cells increased neuronal and BM-associated genes including *Nrn1*, *Nefl*, *Pou4f1*, *Slc17a6* and *Dmrta2*, while GABA^BL^-committed cells increased *Scg2*, *Slc32a1*, *Gad1*, *Gad2* and *Meis2* (Fig. 4e and Supplementary Fig. S13a,b). Fate-probability trajectories showed that both groups retained mixed futures early and separated within a later commitment window (Supplementary Fig. S13c,d).

This axis could be summarized by three branch-control modules. A BM-patterning module contained *Shh*, *Dmrta2* and *Igfbp5*, a BL-selector module contained *Helt*, *Tal2* and *Tle4*, and a BL-reinforcement module contained *Dlk1*, *Lmo1* and *Meis2*. BM-biased founders started higher on the BM-patterning module, BL-biased founders started higher on the BL-selector module, and the BL-reinforcement module rose later along the BL trajectory (Fig. 4f and Supplementary Fig. S12a-c). This ordering separates an early bias in progenitors from a later program that stabilizes the GABA^BL^ outcome.

Finally, in silico perturbation asked how changing each branch program altered the others along the BL trajectory. On BL-committed trajectories, BM-patterning reduced BL-selector activity, while BL-reinforcement reduced BM-patterning and strengthened BL-selector. Later along the same trajectories, BL-reinforcement acquired self-support while retaining its effects on the other two modules (Supplementary Fig. S12d). Together, these analyses support a branch program in which an early patterning-selector imbalance in NPBM progenitors is followed by reinforcement of the committed GABA^BL^ state. This interpretation is consistent with ventral-midbrain biology, where *Shh* and Wnt signalling shape ventral-midbrain patterning and floor-plate neurogenesis^81^, and *Helt* and *Tal2* promote GABAergic fate specification^82,83^. Overall, pandaCREST demonstrates that PhyloFM can convert clone-split ancestry into a continuous hidden-future axis within a progenitor state and connect that axis to interpretable branch programs.

### Branched repair trajectories reconstructed from independently recorded in vivo lineages

Lineage in the zebrafish heart regeneration dataset was recorded using a LINNAEUS-type CRISPR–Cas9 scarring strategy^14,63^. Cas9 and a single-guide RNA targeting the multicopy dTomato transgene of the zebrabow line are injected into one-cell-stage embryos, producing heritable insertions or deletions during early development. After the fish reach adulthood, hearts are cryoinjured and sampled at multiple time points post-injury (0, 3, 7 and 30 dpi in the original study), and single cells are profiled jointly for scar patterns and transcriptome, so that shared scars within a fish reconstruct a lineage tree. Because the scars are generated independently in each embryo, the same scar sequence cannot be treated as a reusable clone label across different fish. PhyloFM therefore compares repair dynamics through the lineage-derived cell-state topology, which provides shared anchors across sampled animals and time points (Fig. 5a). Following Moslin^57^, we restricted our analysis to the first three time points (0, 3 and 7 dpi), which cover the initial injury response and the peak of fibrotic remodelling where the pro-regenerative fibroblast states are most active.

**Figure 5:**
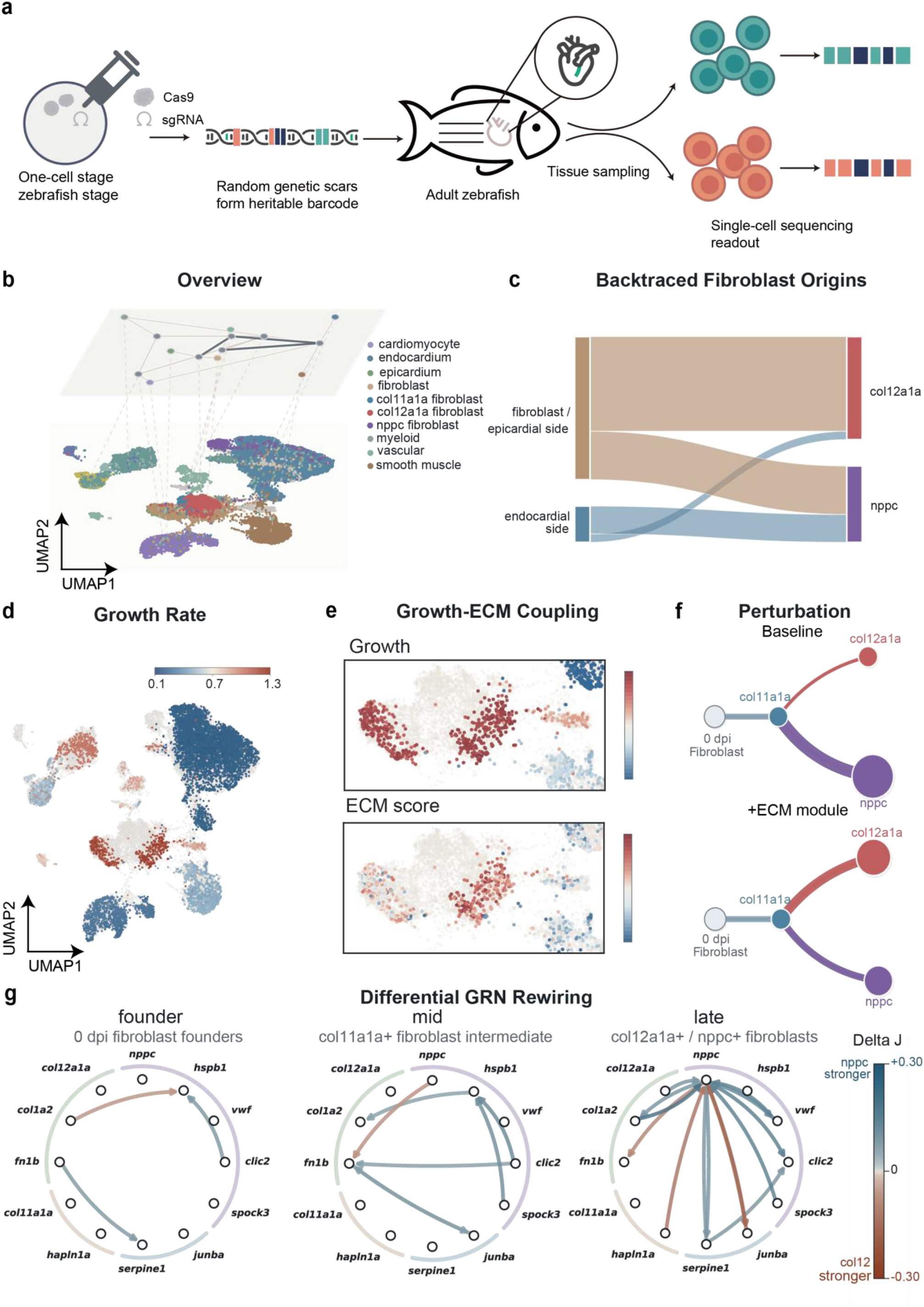
Lineage-aware modelling resolves branched fibroblast repair dynamics in zebrafish heart regeneration. a, LINNAEUS-style CRISPR scarring generates heritable barcodes during early zebrafish development, which are read out with single-cell transcriptomes after adult heart injury. b, Inferred topology and cell-state landscape for the regeneration dataset. c, Backtraced fibroblast origins show distinct source contributions from fibroblast/epicardial and endocardial sides to *col12a1a* and *nppc* terminal fibroblast outcomes. d, Inferred growth-rate field over the repair landscape. e, Coupling between growth and ECM-support scores highlights regions where proliferative and matrix-associated programs co-vary. f, In silico ECM-support perturbation shifts predicted fate mass between *col12a1a* and *nppc* branches. g, Differential velocity-GRN rewiring across founder, intermediate and late trajectory states reveals changing branch support between *nppc* and *col12a1a* programs.

The analysis focused on the fibroblast compartment, where the inferred topology organised *col11a1a*, *col12a1a* and *nppc* fibroblast states along an injury-response route (Fig. 5b). Backtracing the two terminal fibroblast outcomes revealed distinct origins, with the *col12a1a* arm dominated by fibroblast and epicardial-side sources and the *nppc* arm more endocardial-shifted (Fig. 5c). This matches the lineage interpretation in the original study, which placed *col11a1a* and *col12a1a* fibroblasts on the epicardial or constitutive-fibroblast side and *nppc* fibroblasts closer to the endocardial side^63^. As an independent molecular check, differential expression between the model-defined col12a1a-biased and nppc-biased founders, computed on observed 0 dpi expression, recovered the corresponding transient-fibroblast markers, as 15/16 shared markers were enriched on the expected branch. The source-state grouping therefore reproduced the epicardial-side and endocardial-side identities.

The source pool already contained branch information before terminal identities appeared. Among 361 fibroblast source cells at 0 dpi, 51 were biased toward *col12a1a* and 42 toward *nppc*, with only 4 cells shared between the two groups. The nppc-biased group occupied a distinct region of the founder map, and the contrast between ECM and endocardial-shift programs separated the two groups with an AUC of 0.888 (Supplementary Fig. S14a–c).

Continuous trajectories resolved the *nppc* arm as a delayed branch from a shared remodelling corridor. In col12a1a-biased founders, *col12a1a* probability rose early and remained dominant. In the nppc-biased pool, a col12-like phase appeared first and *nppc* identity emerged afterwards (Supplementary Fig. S14d). Module and gene trajectories matched this sequence. ECM support stayed high along the *col12a1a* route, and the endocardial-shift signal rose after the *nppc* branch had committed (Supplementary Fig. S15). Inferred growth was spatially structured and co-localised with ECM support in fibroblast and epicardial regions (Fig. 5d,e and Supplementary Fig. S17d), a placement consistent with the known activation and proliferation of epicardial-derived fibroblasts during zebrafish heart regeneration^84,85^.

In silico perturbation and regulatory sensitivity analyses refined this picture. Increasing the ECM-support program shifted predicted mass toward the *col12a1a* arm and away from *nppc* (Fig. 5f), consistent with ECM acting as a route stabiliser during branch progression. At the founder state, regulatory differences between the two arms were modest, suggesting that early separation is mainly carried by source-cell bias. Later along the trajectory, the *nppc* arm developed a reinforcing *Nppc* and endocardial program that was robust under founder-cell resampling (Fig. 5g and Supplementary Fig. S16a,b), and per-cell *Nppc* and endocardial support rose from the founder to the late state along this branch while remaining flat along the *col12a1a* one (Supplementary Fig. S16a). The genes inside this late regulatory loop are consistent with documented zebrafish heart biology. *vwf*, a canonical endothelial and endocardial marker, reinforces the endocardial-shifted origin of the arm, and on the opposite ECM side of the network, *hapln1a*, known to mark epicardial cells that guide coronary regrowth and revascularisation during regeneration^86^, marks the *col12a1a* route. The mutual edge between *spock3* and *nppc* in the same circuit suggests that spock3 may be a candidate SPOCK-family ECM-associated regulator.

Beyond the fibroblast compartment, other injury-response cell types also carried structured future variation. Inflammatory microstates differed in how committed their futures were: some retained narrow future distributions while others remained broadly uncertain, and in silico perturbation of *apoeb*- and epdl-associated programs in these states shifted predicted inflammatory-processing routes (Supplementary Fig. S17a–c). Overall, zebrafish heart regeneration shows how independently recorded CRISPR scar lineages can be incorporated through cell-state topology. The model recovers a dominant *col12a1a* ECM-remodelling corridor with epicardial-side origins, a smaller endocardial-shifted *nppc* branch that emerges later, and a source-cell asymmetry that predicts the branch outcome.

### Extension to spatial lineage links future fate to the tissue niche

As single-cell lineage tracing is increasingly combined with spatial measurements, we used eTracer to ask whether PhyloFM extends naturally to this emerging data type. eTracer is a recently developed CRISPR– Cas9 lineage tracer that accumulates edits in the 3’UTRs of highly expressed endogenous genes, so evolving barcodes can be read directly from poly(A)-captured transcripts and lineage, single-cell transcriptome and tissue coordinates can be acquired in the same experiment using sequencing-based spatial transcriptomics^19^. Applied to an EGFR-mutant lung adenocarcinoma model under CD8⁺ T-cell cytotoxicity, it produced three serial spatial sections (T0, T1, T2) with shared-clone truth across time and space (Fig. 6a). This is the type of multimodal input that motivated PhyloFM’s modular design. When spatial coordinates are available, the representation stage adds a within-slice neighbour regularizer on the latent space, and the coupling stage extends the hybrid cost with a spatial distance term, so that tissue geometry enters both the latent geometry and the cross-time coupling (Methods).

**Figure 6:**
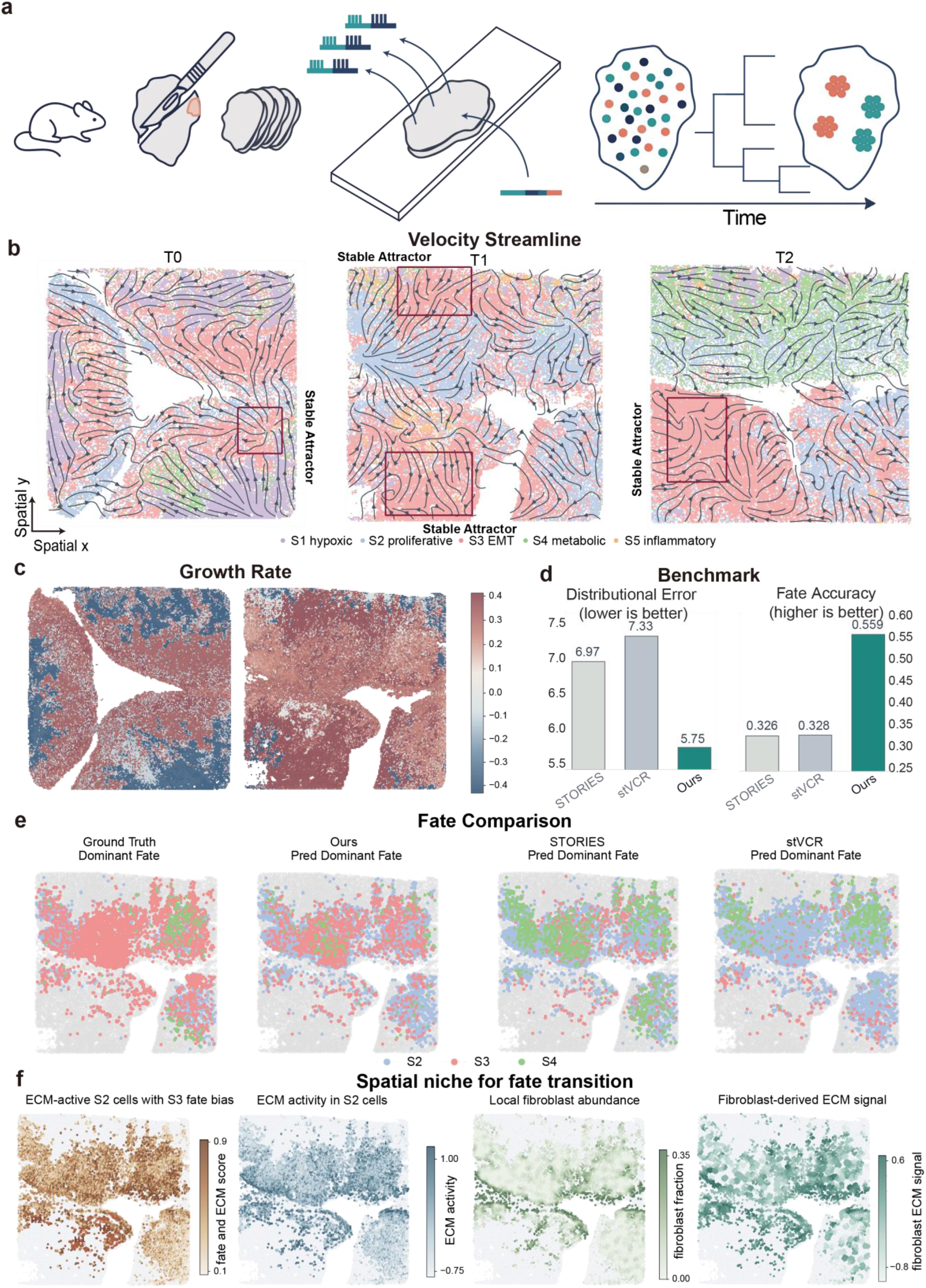
Spatial-lineage modelling of tumour-state progression in eTracer sections. a, eTracer combines static and evolving lineage barcodes, spatial transcriptomics and serial tumour-section sampling to relate clone history, tissue location and transcriptional state. b, PhyloFM velocity streamlines over T0, T1 and T2 sections reveal spatially organized progression through S2 proliferative, S3 EMT-like and related tumour states, including stable attractor regions. c, Inferred spatial growth-rate maps across tumour sections at T0 and T1. d, Benchmark comparison with STORIES and stVCR using distribution error and clone-grounded fate accuracy. e, Dominant-fate maps compare clone-derived ground truth with predictions from PhyloFM, STORIES and stVCR on shared-clone source cells. f, Spatial niche analysis links future-S3-biased ECM-active S2 cells with local fibroblast abundance, ECM activity and fibroblast-derived ECM signalling.

With this spatial extension, PhyloFM remained the most accurate method on the spatiotemporal benchmark, achieving lower W1 than both state-of-the-art spatial baselines STORIES^87^ and stVCR^88^ (5.75 versus 6.97 and 7.33) and higher clone-grounded fate accuracy (0.559 versus 0.326 and 0.328), corresponding to a 17.5% W1 reduction and a 70.4% accuracy gain over the best baseline (Fig. 6d). The recovered dynamics matched the expected tumour-state progression. PhyloFM organized the tumour-progression landscape around an S2 proliferative reservoir feeding a downstream S3 EMT-like state (Fig. 6b), and the inferred transitions supported a directional progression, with most S2 cells remaining S2 but a meaningful fraction moving into S3 between T0 and T1, and S3 becoming the more stable state between T1 and T2 (Supplementary Figs. S18a and S19), in line with the original study^19^. In silico perturbation of S2 programs then showed that ECM- and stress-related programs produced the strongest gains in future-S3 probability, especially in T1 cells with low baseline future-S3 (Supplementary Fig. S18b). Spatial growth maps placed this transition in tissue context, with the strongest mass gain in S2 at T0 shifting toward S3 at later time points (Fig. 6c), and, on T1 shared-clone S2 source cells, PhyloFM reproduced the clone-derived dominant future map more faithfully than either spatial baseline (Fig. 6e).

Because each prediction remains tied to tissue coordinates, the inferred trajectories can be mapped back onto tumour sections. Within T1 S2 cells, regions with high future-S3 probability were concentrated along a restricted tumour interface, where the local ECM signal was high and fibroblasts were both more abundant as neighbours and more active in ECM-related signalling (Fig. 6f). Consistently, genes positively associated with future S3 were enriched for Hallmark EMT, collagen-fibril organization and extracellular-matrix organization (Supplementary Fig. S20), placing the model’s predictions on the expected tumour ECM–EMT axis^89,90^. The eTracer analysis therefore extends the same lineage-topology framework to spatial-lineage data, linking clone-derived future state with tissue niche structure and improving quantitative prediction over spatial baselines.

## Discussion

Lineage-tracing technologies have multiplied over the past decade, yet the data they produce remain difficult to compare. An exact embryonic tree, an inherited barcode, a CRISPR scar, a clone-split label and a spatial clone-family label all report ancestry, but each record supports a different kind of cross-time analysis. The design of PhyloFM follows from this heterogeneity. Cell-state anchors provide the reusable coordinates needed to compare lineage records across time, animals and experimental formats. Topology-regularized representation learning converts those anchors into a latent geometry in which interpolation follows lineage-state geodesics, constraining the learned velocity field along the full transition path as well as at the matched endpoints. Hybrid unbalanced transport then supplies both endpoint matching and a cell-abundance signal for growth learning. When tissue coordinates are measured, spatial regularization and spatial coupling costs extend the same geometry to local tissue structure. The anchor graph becomes the coordinate system on which PhyloFM learns transport, velocity, growth and molecular programs. The anchor graph can come from an exact developmental tree, an inferred differentiation map or another lineage-state graph construction appropriate to the tracing design. A trained model returns population transport, per-cell fate probabilities, commitment timing, growth and module-perturbation responses across lineage technologies.

Across the five benchmarks, lineage information improved performance in every setting we tested, with the form of evidence changing with the lineage record. In *C. elegans,* the exact embryonic tree provided the strongest dynamical benchmark; PhyloFM improved population transport, reduced lineage-local dynamic error and reproduced independently measured EPIC reporter timing for proneural, ciliated-neuron and late-selector programs. In LARRY hematopoiesis, it increased founder-restricted fate accuracy. In pandaCREST, it recovered a clone-aligned future axis within a coarse progenitor state. Ablations that removed or shuffled the lineage signal eliminated these gains. The zebrafish and eTracer analyses extended the same anchor-graph, coupling and velocity-growth design to settings where lineage truth is more indirect, recovering an externally consistent picture of fibroblast remodelling in the regenerating zebrafish heart and transferring to combined spatial-lineage tumour data. Across these scenarios, the shared biological insight is that ancestry reveals future variation before terminal states are evident. PhyloFM turns that signal into continuous trajectories, growth fields and molecular programs, giving heterogeneous lineage records a common dynamical interpretation.

Several limitations and natural extensions remain. First, PhyloFM relies on an informative lineage-state topology. Sparse, noisy or compartment-biased lineage labels will produce a less informative graph, and different tracing formats provide lineage information of different strengths. Second, decoded molecular trajectories are projected gene- or module-level summaries of the learned state representation, while raw gene-expression trajectories are not measured directly. The EPIC comparison in *C. elegans* provides external validation of timing for a subset of reporter genes, but broader gene-level interpretation should be accompanied by readout calibration and, where possible, independent molecular measurements. Third, the inferred growth field is derived from the row mass of the unbalanced OT coupling and reflects an OT-implied mass balance. Direct proliferation or cell-death measurements would be needed to calibrate it as an absolute biological rate. Fourth, module perturbations are model-guided directional steering analyses for hypothesis generation. Experimental perturbation data would be required for causal calibration. Lastly, the spatial term in the hybrid coupling currently uses local Euclidean distance and is best suited to coordinate-aligned tissue sections. A direct extension would be application to the larger spatial-lineage datasets that current technologies are beginning to deliver^19,91^. Further extensions include deeper whole-embryo CRISPR-recorded lineages^15^ and coupling module steering with calibrated single-cell perturbation data to test the directional shifts produced by the model against experimental ground truth^92,93^.

Overall, PhyloFM translates ancestry into a cell-state topology, learns a constrained dynamical geometry and reads out fate, relative cell-abundance change and molecular dynamics from the resulting velocity-growth model. More broadly, PhyloFM moves lineage tracing from a descriptive account of past ancestry toward a unified quantitative framework for reconstructing continuous, interpretable molecular fate dynamics.

## Methods

### Datasets and preprocessing

We analysed five lineage-tracing datasets that span exact embryonic lineage, shared viral barcodes, clone-split CRISPR barcodes, CRISPR scars generated separately in each animal and spatial lineage recording. For each dataset, cells were represented by the first 50 principal components of a log-normalized highly variable gene matrix. When curated PCA coordinates were already provided, they were reused. Otherwise, counts were library-size normalized, log-transformed, filtered to 3,000 highly variable genes by default and projected by PCA with a fixed random seed. The decoder was trained on this representation, and decoded trajectories were mapped back to gene-expression profiles by applying the corresponding PCA basis before summarizing selected genes or gene modules. Lineage records were used to define adjacent-time empirical matchings and a cell-state topology for the representation and coupling steps.

#### *C. elegans* embryonic atlas^62^

We used the lineage-resolved embryonic atlas restricted to ABpxp descendants at 170, 210, 270, 330, 390, 450 and 510 min post-fertilization. This subset contained 6,920 cells in total, with 190, 832, 1,787, 2,192, 855, 640 and 424 cells at the seven stages. The known invariant embryonic lineage supplied the topology directly and was also used for lineage-local dynamic-error evaluation.

#### LARRY hematopoiesis^13^

We used day-2, day-4 and day-6 cells from the raw LARRY hematopoiesis time course, in which shared lentiviral barcodes identify clonal descendants across sampled days. The analysis retained 49,302 cells, with 4,638, 14,985 and 29,679 cells at days 2, 4 and 6. A lineage-positive subset of 33,814 cells was used for representation learning.

#### pandaCREST mouse organoid^17^

We used replicate 1 of the mouse ventral-midbrain clone-split pandaCREST experiment, retaining E11.5 progenitors and day-7 organoid derivatives whose CREST V1 and V2 clone labels were observed at both stages. The analysed subset contained 3,142 cells in total, with 1,173 E11.5 source cells and 1,969 day-7 target cells across 442 shared clones.

#### Zebrafish heart regeneration^63^

We used the lineage-recorded multi-animal zebrafish heart-regeneration series restricted to fibroblast-lineage cells at 0, 3 and 7 dpi. These stages cover the initial injury response and the peak of fibrotic remodelling. The analysed subset contained 44,014 cells in total, with 9,394, 13,783 and 20,837 cells at 0, 3 and 7 dpi.

#### eTracer tumour spatial sections^19^

We used three serial spatial sections of a lineage-traced tumour model, mapped to T0, T1 and T2. The full expression data contained 103,110 cells in total, with 31,953, 35,326 and 35,831 cells across the three sections. For spatial topology learning, 12,000 cells per stage were sampled after clone-internal refinement of the S2 tumour state, while the full PCA representation was retained for dynamical modelling. Spatial coordinates were used as x and y covariates for within-section neighbours and for the spatial term in the hybrid coupling cost.

For the four non-tree datasets, CARTA was used as a standardized topology provider to convert lineage-derived potency or tree summaries into a cell-state graph. Shortest-path distances on this graph were used by the topology loss and lineage-aware coupling. Dataset-specific CARTA inputs, filters and hyperparameters are reported in Supplementary Note S2.

### Lineage-state topology graph

All downstream components of PhyloFM operate on a single structured object: a graph *G* = (*A*, *E*, *w*) over a finite set of cell-state anchors A, with unweighted or lineage-weighted edges E and non-negative edge weights w. Anchors are defined per dataset and include observed cell types and progenitor states from the original study. Each cell i is assigned either a single anchor or a soft anchor distribution *a_i_*, and we write *d_G_*(*a*, *b*) for the shortest-path distance between two anchors.

The graph construction is modular. PhyloFM requires a shortest-path distance on cell-state anchors, and this graph can be supplied by a known developmental tree, an inferred differentiation map, a curated prior or another lineage-graph constructor. For *C. elegans*, where the invariant embryonic lineage is known exactly, we use the published embryonic lineage tree directly. Anchors are cell identities in the tree and *d_G_* is the tree-distance, defined as the number of division edges separating two cells. For the other four datasets, we first reduce the lineage record to the CARTA input format appropriate for that technology. LARRY and pandaCREST are represented as clone-by-terminal-state potency matrices, zebrafish is represented by CRISPR lineage trees with terminal coarse cell-state labels, and eTracer uses a tumour-state CARTA scaffold followed by clone-internal refinement. CARTA returns a binary program matrix whose rows encode progenitor state supports. We convert this matrix into a program graph by immediate subset relations and, in the analyses reported here, attach observed terminal cell types as leaves. We take *d_G_* as the shortest-path distance on this augmented graph. Two cells are called topology-close if *d_G_*(*a_i_*, *a_j_*) is below a soft threshold *d*_0_ and topology-distant otherwise, with the sharpness of this transition controlled by a sharpness hyperparameter introduced in the Topology loss subsection below.

### Topology-regularized latent representation

PhyloFM learns a latent representation with a variational autoencoder. The encoder outputs a Gaussian posterior over each cell, a latent sample is drawn by reparameterization, and the decoder reconstructs the same input feature vector. Both networks are residual MLPs with LayerNorm and SiLU activations. In all analyses reported here, the input feature vector is the 50-dimensional PCA representation used for each adjacent-time comparison, so the reconstruction term is evaluated in PCA space.

For adjacent time points t_k_ and t_k+1_, let (i, j) index a pair of cells sampled from these two time points, with latent codes z_i_, z_j_. The total representation loss is

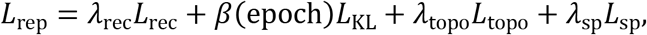

where the reconstruction, KL, topology and (when applicable) spatial-regularization terms are defined below. The spatial term applies only to datasets that come with within-slice neighbour graphs.

#### Reconstruction loss

For a mini-batch B of cells, L_rec_ is the mean squared error between the input expression features and their decoded counterparts:

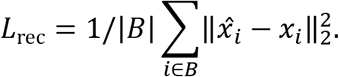

#### KL term

L_KL_ is the standard KL divergence between the posterior and the standard normal prior, averaged over the mini-batch,

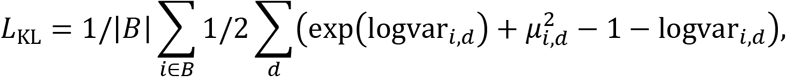

where the KL-divergence weight in the total loss above is annealed under the warm-up schedule *β*(epoch) = *β_start_* + (epoch/epoch_anneal_) × (*β*_max_ − *β*_start_) clamped to *β*_max_.

#### Topology loss

For a sampled pair (i, j) from adjacent time points, write *d_ij_* = *d_G_*(*a_i_*, *a_j_*) and *dz_ij_* = ‖*z_i_* − *z_j_*‖. We turn the graph distance into a soft positive weight

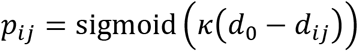

and penalize topology-close pairs with a quadratic attractive term and topology-distant pairs with a hinge-based repulsive term up to a margin γ:

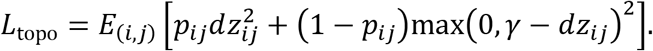

The margin is set analytically from the latent dimensionality as *γ* = sqrt(2 × *d_z_*), so it matches the expected Euclidean distance between two independent draws from the unit Gaussian prior and does not need to be tuned. The two remaining hyperparameters are the soft threshold *d*_0_, which defines the graph distance below which a pair is considered a positive, and the sharpness *κ*, which controls how quickly the soft weight saturates around *d*_0_. The expectation is estimated by random sampling of pairs at each training step.

#### Spatial regularizer (eTracer only)

When within-slice neighbour graphs are available, L_sp_ is a weighted Laplacian smoothness term on the latent space,

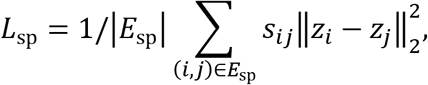

with *s_ij_* the spatial-neighbour edge weights. Setting *λ*_sp_ = 0 recovers the non-spatial model exactly.

The full loss L_rep_ is minimized jointly over φ and θ by Adam with cosine-annealed learning rate.

### Hybrid optimal-transport coupling

For each adjacent gap (*t_k_*, *t*_(*k*+1)_) with *n_0_* source cells and *n*_1_ target cells, we compute a coupling matrix 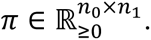 Let *M^expr^* and *M*^lat^ be the pairwise squared Euclidean cost matrices in expression space and in the trained latent space, and *M^sp^* the pairwise squared Euclidean cost in tissue coordinates when spatial data are available. Each cost is mean -normalized and combined into a hybrid cost

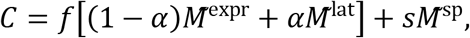

where *α* ∈ [0,1] controls the expression-versus-latent balance, and f and s are feature and spatial weights. Non-spatial datasets set *s* = 0; the feature-versus-spatial balance used for eTracer is (*f*, *s*) = (0.7,0.3).

In the unbalanced mode, π minimizes ⟨*C*, *π*⟩ + *ηH*(*π*) + reg_m_[KL(*π*1‖*μ*) + KL(*π*^T^1‖*ν*)], accommodating mass changes arising from proliferation, death or sampling shifts through a soft marginal relaxation controlled by *reg_m_*. For large datasets, we partition the source and target populations into mini-blocks and solve an unbalanced problem within each block. The block couplings are multiplied by a global factor so that the summed coupling mass matches the target cell count, making row masses comparable across mini-blocks. This block-sparse variant is used for LARRY, pandaCREST, zebrafish and eTracer. In all cases we retain both the coupling *π_ij_* and the source row mass *m_i_* = ∑*_j_ π_ij_* for the next stage.

### Velocity and growth field training

The coupling is used as a training distribution for two neural fields: a latent velocity field *v_ψ_*(*z*, *t*) and a log-growth field *g_ξ_*(*z*, *t*). For a sampled gap k, we draw a coupled pair (*z*_0_, *z*_1_) with probability proportional to *π_ij_* (restricted to source rows with non-negligible mass) and a continuous interpolation parameter *τ* ∼ *U*(0,1), and form the straight latent bridge and its target velocity

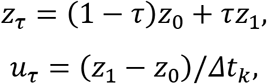

where *Δt_k_* = *t*_(*k*+1)_ − *t_k_* is the physical gap length. The absolute model time is *t_τ_* = *t_k_* + *τ* × *Δt_k_*.

The growth target comes from the coupling row mass of the sampled source cell. For a sampled row i, m_i denotes the row mass and a_i denotes the source reference mass in the same returned-coupling units. We define

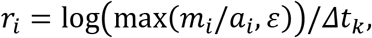

Here *r_i_* is the local log-growth rate relative to the source reference mass. In the count-scaled analyses, *a_i_* is one source-cell unit after coupling rescaling, so this expression is equivalent to *log*(*max*(*m_i_*, *ε*)) / *Δt_k_*.

Both fields are trained jointly by a weighted flow-matching loss

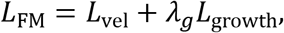

With

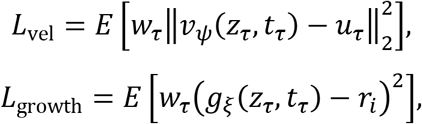

and optional per-sample weight *w_τ_* = *m* that emphasizes low-mass cells near *τ* = 0 and high-mass cells near *τ* = 1; setting *w_τ_* = 1 recovers unweighted flow matching. The expectation runs over gap indices k (sampled uniformly across all experimental gaps in every step), over coupled pairs (*z*_0_, *z*_1_) ∼ *π_k_*, and over *τ* ∼ *U*(0,1), so a single pair of networks (*v_ψ_*, *g_ξ_*) covers the full experimental window.

Training proceeds in three sequential stages. (i) The representation stage minimizes L_rep_ over the encoder φ and decoder θ. (ii) Once trained, (φ, θ) are frozen, and hybrid optimal-transport couplings are computed on the resulting latent codes. (iii) Finally, the velocity field v_ψ_ and growth field g_ξ_ are fitted by minimizing L_FM_, with the couplings serving as the training distribution. Hyperparameters and training schedules for each dataset are documented in Supplementary Note S3.5.

### Inference and downstream quantities

At inference, latent state and log mass are integrated jointly along the learned dynamics,

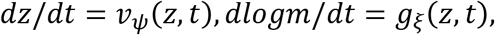

The decoder maps the integrated latent state back to the learned molecular feature space along the trajectory, which is then converted to gene-expression profiles for gene- and module-level analyses. From the same trajectory we extract population predictions used for transport benchmarks, per-cell fate probabilities used for clone-grounded benchmarks and founder-bias maps, commitment positions defined by sustained route-probability support, local growth integrated from the log-mass trajectory, gene-level dynamics, and module-perturbation responses obtained by defining a module direction in gene space and re-integrating after the corresponding displacement.

Per-cell fate probabilities were computed with a shared k-nearest-neighbour procedure on the model state at the evaluation time. For each analysis, observed reference cells were selected from the relevant state-labelled population at the target time or from the observed state-labelled population used for continuous route analyses. After a source cell was integrated and mapped to the learned molecular feature space, its probability for fate class c was the normalized inverse-distance weight assigned to reference neighbours carrying label c, using k = 75 with k capped by the reference population size. Clone-level predictions were obtained by averaging these per-cell probabilities over source cells from the same clone, and clone-derived ground truth was the observed fraction of target descendants from that clone in each fate class.

### Route, branch and molecular analyses

Route and branch analyses were derived from the integrated trajectories and the fate-probability estimates described above. Dataset-specific terminal labels were grouped into the biological routes shown in each figure. For each source cell, the dominant route was the route with the largest terminal probability. Commitment position was taken as the earliest point along the integrated trajectory at which the eventual dominant route reached sustained support for the remainder of the trajectory. Branch and founder cohorts were defined from terminal route probabilities or clone-derived fate margins, and the exact label groupings and cutoffs are provided with the figure source files.

Molecular trajectories were computed on the same integration grid. Decoded PCA states were projected back to selected gene profiles with the dataset-specific PCA basis, and module scores were calculated as signed, standardized averages of the selected genes. Temporal gene or module patterns were assigned from route-mean curves, and differential-expression or enrichment summaries used the observed cells assigned to the corresponding route or branch cohort.

### Benchmarks and ablations

Population transport was evaluated using Wasserstein-1 (W1) and Wasserstein-2 (W2) distances between predicted and observed populations at each time point. For the *C. elegans* time-point hold-out transport interpolation benchmark, each interior stage was omitted in turn. In each fold, the topology-regularized VAE and the velocity and growth fields were retrained without cells from the omitted stage or direct couplings to that stage. Flow matching was trained across the interval between the two flanking observed stages, and trajectories initiated at 170 min were scored against the omitted population. Lineage-local accuracy was evaluated using lineage-local dynamic error in *C. elegans*, which compares one-step model predictions to the next-time population connected by the known tree. For the founder-level fate accuracy in the LARRY dataset, day-2 source cells were propagated to day 6, assigned terminal fates by a shared k-nearest-neighbour vote against observed day-6 cells and scored against clone-derived fate labels. Hidden-future accuracy in pandaCREST was evaluated as Spearman and Pearson correlations between the latent GLUBM minus GABABL fate margin and the empirical clone-derived margin on NPBM cells. Clone-grounded fate accuracy in eTracer was evaluated on T1 shared-clone S2 source cells. Recovery of documented regulatory relationships in C. elegans and LARRY was evaluated using conditional mutual information to assess directionality in generated trajectories, using the relationship panels described in the Supplementary Notes. Baselines comprised VGFM, TIGON, scDiffEq and OT-CFM for non-spatial datasets, and STORIES and stVCR for the spatial eTracer benchmark.

Ablation experiments removed or perturbed specific components of the lineage signal: (i) removing the topology graph entirely, (ii) shuffling lineage labels, (iii) reweighting the topology and reconstruction terms in the representation loss, and (iv) substituting a PCA representation for the topology-regularized latent space. All ablations were trained and evaluated on identical splits, allowing differences in transport, fate or local dynamic metrics to be assessed with respect to the manipulated component.

### Module-perturbation analyses

A gene module M is a set of marker or regulator genes together with per-gene signs. Given a cell i with expression x_i_, we compute a module direction in gene space as the signed z-scored mean of the module genes, and project it into the latent space via the encoder Jacobian at the source cell,

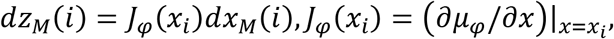

where *dx_M_*(*i*) is the gene-space direction. A perturbation is applied by displacing the source latent code along *dz_M_*(*i*) by a scaled step 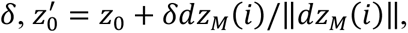 and re-integrating the learned dynamics from 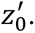 The perturbation magnitude δ is chosen per module to remain within the empirical latent-space distribution (below the median nearest-neighbour distance in latent space). Responses are summarized as shifts in dominant terminal fate fractions, cohort-level fate margins or route-specific module scores.

### Local influence and spatial-neighbourhood analyses

Velocity-network and gene-regulatory rewiring panels were sensitivity summaries of the trained velocity field. At representative founder, intermediate and late trajectory states, small signed gene or module displacements were applied in PCA feature space and the change in the next-step decoded velocity was projected onto selected output genes or modules. Influence matrices were averaged over cells within each route state, and branch-differential edges were reported as differences between matched route states. Spatial-neighbourhood analyses in eTracer used tissue-coordinate nearest neighbours to compare local cell-type fractions, growth, AP-1 score and predicted future-state probability between high- and low-probability S2 cells, followed by regression and Moran’s I summaries.

### Cell-state Sankey diagrams

Sankey diagrams of cell-state transitions were constructed from integrated trajectories by binning cells at successive integration times and computing the empirical transition probabilities across binned cell-state labels at consecutive time bins. The resulting flows visualize how the model redistributes mass between cell-state categories under its learned dynamics, complementing the per-cell fate probabilities defined in Inference and downstream quantities. Trajectory assignments and transition diagnostics are reported in the corresponding supplementary source tables.

## Code and data availability

PhyloFM is implemented in PyTorch and is available at https://github.com/JackkWangzh/PhyloFM. Preprocessing scripts, anchor-graph construction, model checkpoints and figure-generation scripts for each dataset will be released alongside the paper, with traceability files linking every main and supplementary panel to the script that produced it. Public datasets were obtained from the references cited in the Datasets and preprocessing section.

## Acknowledgements

X.Y. and C.T. acknowledge support from the National Natural Science Foundation of China (NSFC; grant nos. 92570207 and 32088101) and the Fundamental and Interdisciplinary Disciplines Breakthrough Plan of the Ministry of Education of China (grant no. JYB2025XDXM502). P.Z. acknowledges support from the NSFC (grant nos. 12288101, 8206100646, and T2321001), the Fundamental and Interdisciplinary Disciplines Breakthrough Plan of the Ministry of Education of China (grant no. JYB2025XDXM502), and the Fundamental Research Funds for the Central Universities. We also acknowledge the High-performance Computing Platform of Peking University for providing computational resources.

## Author contributions

P.Z., X.Y., C.T., and Z.W. conceived this research. Z.W. and Z.Z. designed the algorithm. Z.W. analysed the data and prepared the visualizations. All authors interpreted the results. Z.W. and Z.Z. wrote the original draft. P.Z., X.Y., and C.T. reviewed and edited the manuscript, and supervised the work. All authors read and approved the manuscript.

