## Supplementary Information for "Reconstructing lineage-constrained gene-program dynamics with PhyloFM"

### Supplementary Information for
Reconstructing lineage-constrained gene-program dynamics with PhyloFM

#### Contents

**Supplementary Note S1. Transport background for time-resolved single-cell dynamics** **3**

**Supplementary Note S2. Datasets, preprocessing and lineage-state topology** **5**

**Supplementary Note S3. PhyloFM training, inference and downstream analyses** **6**

**Supplementary Note S4. Benchmark design, baselines and metrics** **17**

**Supplementary Note S5. Lineage-record formats and anchor-topology rationale** **19**

**Supplementary Note S6. Capability matrix of PhyloFM and related methods** **21**

**Supplementary Figure S1. Benchmark robustness and lineage-ablation analysis in *C. elegans*** **23**

**Supplementary Figure S2. Lineage-aware Moslin-FM and static-coupling comparisons in *C. elegans*** **24**

**Supplementary Figure S3. Inferred fine cell-state flow across *C. elegans* neurodevelopment** **25**

**Supplementary Figure S4. Module ontology support, founder marker maps and route-specific molecular dynamics** **26**

**Supplementary Figure S5. Documented regulatory relationships in *C. elegans* neuronal trajectories** **27**

**Supplementary Figure S6. EPIC GFP reporter validation of inferred *C. elegans* gene dynamics** **28**

**Supplementary Figure S7. Comparative founder fate maps and accuracy in LARRY hematopoiesis** **29**

**Supplementary Figure S8. Growth-associated structure in LARRY hematopoiesis** **30**

**Supplementary Figure S9. Molecular support for hidden founder fate and branch dynamics** **31**

**Supplementary Figure S10. Documented myeloid regulatory relationships in LARRY hematopoiesis** **32**

**Supplementary Figure S11. Branch-driver support for pandaCREST NPBM fate structure** **33**

**Supplementary Figure S12. Three-module support for the NPBM branch-control model** **34**

**Supplementary Figure S13. Gene-program validation and timing of NPBM branch commitment** **35**

**Supplementary Figure S14. Founder bias and timing of the zebrafish fibroblast branch** **36**

**Supplementary Figure S15. Gene-program timing along zebrafish fibroblast routes** **37**

**Supplementary Figure S16. Velocity-GRN support for late *nppc* self-reinforcement** **38**

**Supplementary Figure S17. Secondary plasticity, inflammatory microstate and growth-ECM support analyses** **39**

**Supplementary Figure S18. Tumour-state transitions and module perturbation support in eTracer** **40**

**Supplementary Figure S19. Continuous tumour-state composition across eTracer progression** **41**

**Supplementary Figure S20. Functional enrichment of future-S3-associated genes** **42**

### Supplementary Note S1. Transport background for time-resolved single-cell dynamics

#### S1.1 From sampled cell states to population transport

Single-cell time series measure different cells at different experimental times. The basic mathematical object is therefore a sequence of empirical population measures. If cells at time $t_k$ have state vectors $\mathbf{x}_i^{(k)} \in \mathbb{R}^d$ and optional weights $a_i^{(k)}$, the observed population is

$$\mu_k = \sum_{i=1}^{n_k} a_i^{(k)} \delta_{\mathbf{x}_i^{(k)}}, \qquad a_i^{(k)} \geq 0. \tag{S1}$$

The state vector can be a transcriptomic embedding, a learned latent state or a joint transcriptomic-spatial representation. Trajectory inference from time-series single-cell data asks how mass should move from $\mu_k$ to $\mu_{k+1}$ and how this movement should be interpolated between observed times. Optimal transport (OT) has become a standard language for this problem because it links unpaired populations through an explicit transport cost [1, 2].

#### S1.2 Static and dynamical optimal transport

For two adjacent times, static OT searches for a nonnegative coupling matrix $\pi \in \mathbb{R}_+^{n_k \times n_{k+1}}$ that transports mass from the source cells to the target cells at minimal cost [3]. In the balanced case,

$$\min_{\pi \geq 0} \sum_{i,j} \pi_{ij} c(\mathbf{x}_i^{(k)}, \mathbf{x}_j^{(k+1)}) \quad \text{s.t.} \quad \sum_j \pi_{ij} = a_i^{(k)}, \quad \sum_i \pi_{ij} = a_j^{(k+1)}. \tag{S2}$$

The cost $c$ usually measures transcriptomic, latent, spatial or combined dissimilarity. In practice, problem (S2) is regularized by adding an entropy term $-\varepsilon H(\pi) = \varepsilon \sum_{ij} \pi_{ij}(\log \pi_{ij} - 1)$. This makes the objective strictly convex and permits solution by Sinkhorn iteration [4]. The regularization strength $\varepsilon$ controls the entropy of the resulting plan. Static OT estimates correspondences between adjacent time points. Continuous interpolation requires an additional dynamical model.

Dynamical OT writes the same problem as a continuous population flow. A density $\rho_t$ and velocity field $\mathbf{v}_t$ satisfy the continuity equation

$$\partial_t \rho_t(\mathbf{x}) + \nabla \cdot \big(\rho_t(\mathbf{x}) \mathbf{v}_t(\mathbf{x})\big) = 0, \tag{S3}$$

and the Benamou-Brenier formulation minimizes kinetic action [5],

$$\mathcal{A}_{\mathrm{OT}}(\rho, \mathbf{v}) = \int_0^1 \int_{\mathbb{R}^d} \frac{1}{2} \|\mathbf{v}_t(\mathbf{x})\|^2 \rho_t(\mathbf{x}) \, \mathrm{d}\mathbf{x} \, \mathrm{d}t. \tag{S4}$$

Neural dynamical OT methods use this view to learn continuous cell-state flows from discrete time points [6]. The mass-preserving assumption is restrictive for developmental and disease processes because proliferation, death, differentiation-associated expansion and sampling changes alter total mass.

#### S1.3 Unbalanced optimal transport and growth

Unbalanced optimal transport relaxes the marginal constraints so that source and target masses can differ [7–9]. A common static formulation penalizes deviation from the observed marginals with divergences,

$$\min_{\pi \geq 0} \sum_{i,j} \pi_{ij} c_{ij} \; + \; \lambda_0 D\Big(\pi \mathbf{1} \,\Big\|\, a^{(k)}\Big) \; + \; \lambda_1 D\Big(\pi^\top \mathbf{1} \,\Big\|\, a^{(k+1)}\Big), \tag{S5}$$

where $D$ is often a KL-type divergence. The row sum of $\pi$ then gives the amount of source mass used by the coupling, and the column sum gives the amount of target mass explained by the coupling.

The corresponding dynamical formulation adds a local source term $g_t$,

$$\partial_t \rho_t(\mathbf{x}) + \nabla \cdot \big(\rho_t(\mathbf{x})\mathbf{v}_t(\mathbf{x})\big) = g_t(\mathbf{x})\rho_t(\mathbf{x}). \tag{S6}$$

Here $g_t > 0$ denotes local expansion and $g_t < 0$ denotes local depletion. Together with a Fisher–Rao growth penalty proportional to $\int_0^1\!\int g_t^2 \rho_t \,\mathrm{d}\mathbf{x}\,\mathrm{d}t$, this source-augmented continuity equation gives the Wasserstein–Fisher–Rao, or Hellinger–Kantorovich, formulation of unbalanced dynamical transport [8, 9]. Recent single-cell methods use this view to model net population expansion or depletion together with state transition, including regularized unbalanced OT and velocity-growth models [10–13]. In practice, an adjacent-time coupling can provide both transition targets and a mass signal. The couplings used here are scaled to the observed target population size, with each source cell assigned unit initial mass. For a source cell $i$, the row mass

$$m_i = \sum_j \pi_{ij} \tag{S7}$$

is converted into a local log-mass-change target,

$$r_i = \frac{\log(\max(m_i, \epsilon))}{\Delta\tau}. \tag{S8}$$

Here $\Delta\tau$ is the normalized model-time interval between adjacent sampled snapshots. The learned field describes the net population-mass change implied by the fitted unbalanced coupling over that interval.

#### S1.4 Flow matching

Flow matching learns a time-dependent vector field by regressing velocities along conditional paths [14, 15]. Given paired endpoints $(\mathbf{z}_0, \mathbf{z}_1)$ in a representation space and an interpolation time $\tau \sim \mathrm{Unif}(0, 1)$, a simple conditional path is

$$\mathbf{z}_\tau = (1 - \tau)\mathbf{z}_0 + \tau\mathbf{z}_1. \tag{S9}$$

The target velocity along this path is

$$\mathbf{u}_\tau = \mathbf{z}_1 - \mathbf{z}_0. \tag{S10}$$

A neural field $v_\theta(\mathbf{z}, \tau)$ is trained with

$$\mathcal{L}_{\mathrm{FM}}(\theta) = \mathbb{E}_{\mathbf{z}_0,\mathbf{z}_1,\tau}\left[\|v_\theta(\mathbf{z}_\tau, \tau) - (\mathbf{z}_1 - \mathbf{z}_0)\|^2\right]. \tag{S11}$$

More general paths, stochastic bridges and score-flow formulations connect flow matching to Schrödinger bridge learning [16]. In single-cell modeling, flow-based generative approaches have been applied to temporal population dynamics and perturbation-conditioned phenotype prediction [12, 17, 18]. Its practical appeal is that endpoint couplings can be reused as supervised training pairs for a continuous velocity field.

#### Supplementary Note S2. Datasets, preprocessing and lineage-state topology

This note describes expression preprocessing, lineage-topology construction and the cell populations analysed with PhyloFM. The invariant embryonic tree is used directly for *C. elegans*, and CARTA provides a differentiation graph for the other datasets. PhyloFM combines PCA representations with distances on these lineage-derived graphs. Couplings between adjacent sampled times are then estimated from molecular, latent and, where available, spatial distances.

**Expression preprocessing and adjacent-time input construction.** All analyses use 50-dimensional PCA features as model input. Existing PCA representations were retained when available. Otherwise, expression counts were library-size normalized and log transformed, highly variable genes were selected using a default target of 3,000 genes, and PCA was performed with a fixed random seed. Lineage records define the cell-state topology and provide clone-grounded targets for evaluation when shared clones connect sampled times. The eTracer analysis additionally incorporated local spatial neighbourhoods within each section.

The decoder reconstructs the same PCA representation used for adjacent-time comparisons. For gene-level and module-level analyses, decoded PCA trajectories are projected back to the highly variable gene-expression space using the dataset-specific PCA basis and preprocessing mean. Gene and module scores are then calculated from these reconstructed expression values. Module perturbations follow the corresponding gene-module directions in the learned latent space.

**CARTA-derived differentiation graph.** CARTA estimates a differentiation map represented as a graph. Internal nodes correspond to progenitor programs defined by the terminal fates they support, and edges connect programs with successive levels of fate restriction. Observed terminal cell types or states are connected to the corresponding progenitor nodes, while undifferentiated cells remain associated with internal nodes. PhyloFM calculates shortest-path distances on the undirected form of this graph. For cells $i$ and $j$ assigned to anchors $a_i$ and $a_j$, the topology dissimilarity used during representation learning is

$$d_{\mathrm{topo}}(i, j) = 2\{d_{\mathrm{sp}}(a_i, a_j) + 1\},$$

where $d_{\mathrm{sp}}$ is the shortest-path distance between the assigned anchors on the differentiation graph. This graph-derived dissimilarity shapes the latent representation used to estimate adjacent-time couplings.

**Dataset-specific lineage topology and analysed populations.** Cell counts give the populations included in each analysis.

| Dataset | Lineage information | Differentiation topology | Analysed data |
|---|---|---|---|
| *C. elegans* | Published invariant ABpxp embryonic tree | The known developmental tree is used directly | 6,920 cells across seven stages at 170, 210, 270, 330, 390, 450 and 510 min |
| LARRY | Shared lentiviral clones linking day-2 progenitors to their later descendants | CARTA inferred a graph with ten progenitor programs and nine terminal fates from clone-level fate potency | 49,302 cells across days 2, 4 and 6. The lineage-positive subset used for representation learning contained 33,814 cells |
| panda-CREST | Shared CREST clone labels linking E11.5 progenitors and day-7 organoid descendants | CARTA inferred six progenitor programs connected to 12 differentiated terminal states | 3,142 cells, comprising 1,173 E11.5 cells and 1,969 day-7 cells across 442 shared clones |
| Zebrafish | CRISPR lineage trees with terminal cells annotated by 14 cardiac cell states | CARTA integrated the lineage trees into a graph with seven progenitor programs and 14 terminal cardiac states | 44,014 fibroblast-lineage cells across 0, 3 and 7 dpi |
| eTracer | Spatial clone records with tumour and microenvironment state annotations | A CARTA-derived tumour-state graph informed by clone-level state composition was combined with local spatial neighbourhoods | 103,110 cells across three sections. Topology learning used 12,000 cells per stage, and dynamical modelling retained the full PCA representation |

**Dataset-specific CARTA construction.** For LARRY and pandaCREST, clone-level fate potency was used to infer progenitor relationships and their associated terminal states. For zebrafish heart regeneration, CARTA integrated CRISPR lineage trees whose terminal cells were mapped to 14 cardiac states. For eTracer, the tumour-state graph incorporated clone-level state composition across adjacent sections and was combined with within-section spatial neighbourhoods. These graphs define the cell-state anchors used by PhyloFM.

### Supplementary Note S3. PhyloFM training, inference and downstream analyses

PhyloFM is trained in four sequential stages: (i) construction of a lineage-shaped cell-state graph, (ii) representation learning that maps cells into a latent space whose pairwise geometry is shaped by the graph, (iii) computation of one hybrid unbalanced OT coupling per adjacent time gap on the trained latent codes, and (iv) simulation-free flow-matching training of a latent velocity field and a growth field on those couplings. After training, latent state and log-mass are integrated jointly through the velocity and growth fields. The decoder reconstructs the PCA representation, which is then projected back to gene-expression space with the dataset-specific PCA basis for gene-level and module-level analyses. This note formalizes the training procedure as pseudocode (Algorithms S1–S3) and lists the per-dataset hyperparameters used in each analysis (Tables S1–S5).

#### S3.1 Notation

- $\mathbf{x}_i^{(k)} \in \mathbb{R}^{d_x}$: PCA feature vector of the $i$-th cell at snapshot $k$. In the analyses reported here $d_x = 50$. The decoder reconstructs this representation, and gene values are obtained through the dataset-specific PCA basis.
- $\ell_i$: lineage record (tree edge, clone label, barcode, or scar pattern).
- $\mathbf{s}_i^{(k)} \in \mathbb{R}^2$: tissue coordinates when spatial data are available.
- $G = (\mathcal{A}, \mathcal{E}, w)$: lineage-shaped anchor graph on cell-state anchors $\mathcal{A}$; $D_{ij}$ the graph-shortest-path distance pulled back to cells $(i, j)$.

- $\mathbf{z}_i \in \mathbb{R}^{d_z}$: latent code; $\phi$, $\theta$: encoder and decoder networks.
- $\mathbf{v}_\psi : \mathbb{R}^{d_z} \times [0,T] \to \mathbb{R}^{d_z}$: latent velocity field. $g_\xi : \mathbb{R}^{d_z} \times [0,T] \to \mathbb{R}$: latent log-mass-change field.
- $\pi_k \in \mathbb{R}_+^{n_k \times n_{k+1}}$: hybrid OT coupling between snapshots $k$ and $k$+1; $m_k(i) = \sum_j \pi_k(i,j)$ is its source row mass relative to unit initial mass for source cell $i$.
- $\Delta\tau_k$: normalized model-time interval between snapshots $k$ and $k$+1; adjacent intervals have unit duration.
- $\alpha, f, s$: latent-vs-feature, feature-coupling and spatial-coupling weights in the cost. $\varepsilon$, $\rho$: entropy and marginal regularizations of the unbalanced OT solver. $\lambda_{\text{rec}}, \lambda_{\text{top}}, \lambda_{\text{sp}}, \lambda_g$: loss weights. $\beta_{\max}$: maximum KL weight after warm-up.

#### S3.2 Training pipeline

The full training procedure is summarized in Algorithm S1. Stage 1 turns lineage records into a graph distance over cells, against which the encoder is shaped in Stage 2. Stage 3 then computes one hybrid OT coupling per adjacent gap on the resulting latent codes, by calling the subroutine in Algorithm S2. Stage 4 fits the velocity and growth fields on these couplings.

For a sampled pair of cells $(i,j)$, let $d^z_{ij} = \|\mathbf{z}_i - \mathbf{z}_j\|_2$ and let $D_{ij}$ be their distance on the lineage-derived graph. The topology weight

$$w_{ij} = \sigma[\kappa(\tau - D_{ij})] \tag{S12}$$

varies smoothly from nearby to distant lineage relationships. The topology loss is

$$L_{\text{top}} = \frac{1}{|B|} \sum_{(i,j)\in B} \Big[ w_{ij}(d^z_{ij})^2 + (1 - w_{ij}) \big[M - d^z_{ij}\big]_+^2 \Big], \tag{S13}$$

which brings lineage-near cells together and separates lineage-distant cells by margin $M$. All analyses used $\tau = 4$, $\kappa = 1$ and $M = \sqrt{2d_z}$.

**Algorithm S1** PhyloFM training pipeline

**Require:** Snapshots $\{(\mathbf{x}_i^{(k)}, \ell_i)\}_{k=0}^{T}$; optional spatial coordinates $\{\mathbf{s}_i^{(k)}\}$
**Require:** Hyperparameters $\lambda_{\text{rec}}, \lambda_{\text{top}}, \lambda_{\text{sp}}, \beta_{\max}$; $\alpha, f, s, \varepsilon, \rho, \lambda_g$; epoch counts $E_{\text{VAE}}, E_{\text{FM}}$
**Ensure:** Trained encoder $\phi$, decoder $\theta$, velocity $\mathbf{v}_\psi$, growth $g_\xi$

**Stage 1.** Assign cells to anchors from the published lineage tree or CARTA-derived differentiation graph $G = (\mathcal{A}, \mathcal{E}, w)$.
Compute pairwise graph-shortest-path distance $D_{ij}$ between the anchors assigned to cells $i$ and $j$.

**Stage 2.** Initialize $\phi, \theta$.
**for** epoch $= 1, \dots, E_{\text{VAE}}$ **do**
  $\beta \leftarrow \min\big(\frac{\text{epoch}}{E_{\text{VAE}}}, 1\big) \cdot \beta_{\max}$ ▷ linear KL warm-up
  Sample cross-time cell pairs $\{(i, j)\}$ and retrieve their graph distances $D_{ij}$.
  $(\boldsymbol{\mu}, \log \boldsymbol{\sigma}^2) \leftarrow \phi(\mathbf{x}_i)$; $\mathbf{z}_i \leftarrow \boldsymbol{\mu} + \boldsymbol{\sigma} \odot \boldsymbol{\epsilon}$, $\boldsymbol{\epsilon} \sim \mathcal{N}(\mathbf{0}, \mathbf{I})$ (analogously for $j$).
  $\hat{\mathbf{x}} \leftarrow \theta(\mathbf{z})$.
  $L_{\text{rec}} \leftarrow \frac{1}{|B|} \sum \|\hat{\mathbf{x}} - \mathbf{x}\|_2^2$
  Compute $L_{\text{top}}$ from Eq. S13.
  $L_{\text{KL}} \leftarrow \frac{1}{|B|} \sum \text{KL}\big(\mathcal{N}(\boldsymbol{\mu}, \boldsymbol{\sigma}^2) \,\|\, \mathcal{N}(\mathbf{0}, \mathbf{I})\big)$
  **if** within-slice spatial graph available **then**
    $L_{\text{sp}} \leftarrow \frac{1}{|E_{\text{sp}}|} \sum_{(u,v) \in E_{\text{sp}}} w_{uv} \|\mathbf{z}_u - \mathbf{z}_v\|_2^2$ ▷ $k$-NN Laplacian on tissue coordinates
  **else**
    $L_{\text{sp}} \leftarrow 0$
  **end if**
  $L \leftarrow \lambda_{\text{rec}} L_{\text{rec}} + \lambda_{\text{top}} L_{\text{top}} + \beta L_{\text{KL}} + \lambda_{\text{sp}} L_{\text{sp}}$
  Adam step on $(\phi, \theta)$ with $\nabla L$.
**end for**

**Stage 3.** Freeze $(\phi, \theta)$; set $\mathbf{z}_i^{(k)} \leftarrow \phi(\mathbf{x}_i^{(k)})$ for all $i, k$.
**for** $k = 0, \dots, T-1$ **do**
  $\pi_k \leftarrow \text{HybridUOT}\big(\mathbf{x}^{(k)}, \mathbf{x}^{(k+1)}, \mathbf{z}^{(k)}, \mathbf{z}^{(k+1)}, \mathbf{s}^{(k)}, \mathbf{s}^{(k+1)}; \alpha, f, s, \varepsilon, \rho\big)$
  $m_k(i) \leftarrow \sum_j \pi_k(i, j)$.
**end for**

**Stage 4.** Initialize $\mathbf{v}_\psi, g_\xi$.
**for** epoch $= 1, \dots, E_{\text{FM}}$ **do**
  Sample gap $k$; sample $(i, j) \sim \pi_k / \sum \pi_k$; sample $u \sim \text{Uniform}(0, 1)$.
  $t \leftarrow t_k + u \Delta\tau_k$; $\mathbf{z}_t \leftarrow (1-u)\, \mathbf{z}_i^{(k)} + u\, \mathbf{z}_j^{(k+1)}$
  $\mathbf{u} \leftarrow \big(\mathbf{z}_j^{(k+1)} - \mathbf{z}_i^{(k)}\big) / \Delta\tau_k$ ▷ target velocity
  $r \leftarrow \log\big(\max(m_k(i), \epsilon)\big) / \Delta\tau_k$ ▷ target log-mass change
  $L_v \leftarrow \|\mathbf{v}_\psi(\mathbf{z}_t, t) - \mathbf{u}\|_2^2$
  $L_g \leftarrow \big(g_\xi(\mathbf{z}_t, t) - r\big)^2$
  Adam step on $(\psi, \xi)$ with $\nabla(L_v + \lambda_g L_g)$.
**end for**
**return** $(\phi, \theta, \mathbf{v}_\psi, g_\xi)$.

##### S3.3 Hybrid optimal-transport coupling

Stage 3 in Algorithm S1 relies on the subroutine in Algorithm S2, which assembles a hybrid cost and solves an entropy-regularized unbalanced OT problem. The cost combines mean-normalized pairwise distances in PCA space and the topology-shaped latent space. For eTracer, physical distance contributes an additional spatial term.

---

**Algorithm S2** HybridUOT: hybrid unbalanced OT coupling

---

**Require:** Source/target features $(\mathbf{x}^a, \mathbf{x}^b)$, latent codes $(\mathbf{z}^a, \mathbf{z}^b)$, spatial coordinates $(\mathbf{s}^a, \mathbf{s}^b)$ if available
**Require:** Latent-feature mix $\alpha \in [0, 1]$, feature weight $f$, spatial weight $s$ (set $s = 0$ if no spatial), entropy reg. $\varepsilon$, marginal reg. $\rho$
**Ensure:** Coupling $\pi \in \mathbb{R}_+^{n \times m}$

$C^x_{ij} \leftarrow \|\mathbf{x}^a_i - \mathbf{x}^b_j\|_2^2;\quad C^x \leftarrow C^x / \overline{C^x}$
$C^z_{ij} \leftarrow \|\mathbf{z}^a_i - \mathbf{z}^b_j\|_2^2;\quad C^z \leftarrow C^z / \overline{C^z}$
**if** spatial provided **then**
    $C^s_{ij} \leftarrow \|\mathbf{s}^a_i - \mathbf{s}^b_j\|_2^2;\quad C^s \leftarrow C^s / \overline{C^s}$
**else**
    $C^s \leftarrow \mathbf{0}$
**end if**
$C^{\text{feat}} \leftarrow (1 - \alpha)\, C^x + \alpha\, C^z$
$C \leftarrow f\, C^{\text{feat}} + s\, C^s$
$\pi \leftarrow \mathrm{UOT}\big(\mathbf{a}, \mathbf{b}, C; \varepsilon, \rho\big)$.
Normalize $\pi$ to the target population mass used in the analysis.
For blockwise UOT, solve normalized subproblems and apply a common rescaling so that the assembled coupling has the same target population mass.
**return** $\pi$

---

##### S3.4 Inference

At inference (Algorithm S3), the trained velocity and growth networks are integrated forward in latent space from a chosen initial population. The latent trajectories are decoded to PCA space and projected back to gene-expression space for expression-level analyses. The same procedure produces population transport, per-cell fate probabilities, in-silico module perturbations and continuous gene trajectories used in the main text.

Per-cell fate probabilities are computed by a shared $k$-nearest-neighbour state-assignment procedure on the model state at the evaluation time. For a decoded query state $\hat{\mathbf{x}}_i(t)$, let $\mathcal{N}_k(i)$ be its $k$ nearest observed reference cells in the relevant target or state-labelled reference population, with reference labels $y_q$ and distances $d_{iq}$. The procedure uses inverse-distance weights

$$w_{iq} = \frac{(d_{iq} + 10^{-6})^{-1}}{\sum_{q' \in \mathcal{N}_k(i)} (d_{iq'} + 10^{-6})^{-1}}, \tag{S14}$$

and assigns fate probability

$$p_i(c \mid t) = \sum_{q \in \mathcal{N}_k(i)} w_{iq}\, \mathbf{1}\{y_q = c\}. \tag{S15}$$

Unless stated otherwise for a specific benchmark or the reference set is smaller, $k = 75$. Clone-level predictions are obtained by averaging $p_i(c \mid t)$ over source cells from the same clone. Clone-grounded target probabilities are empirical descendant fractions in the target population, computed from the same barcode or clone label.

Route and branch probabilities are obtained by summing fate probabilities over figure-specific terminal label groups. If $\mathcal{R}$ is a set of terminal labels defining a biological route, then

$$p_i(\mathcal{R} \mid t) = \sum_{c \in \mathcal{R}} p_i(c \mid t). \tag{S16}$$

The dominant route of a source cell is the route with the largest terminal value of $p_i(\mathcal{R} \mid t)$. Commitment time is the earliest integration-grid point from which support for the eventual dominant route is maintained through the remainder of the trajectory. Route definitions and support thresholds were specified for each dataset according to its terminal cell-state annotations.

Gene and module trajectories are computed on the same integration grid as the fate probabilities. A decoded PCA trajectory $\hat{\mathbf{x}}_i(t)$ is projected to selected genes using the PCA basis for that dataset. For a signed module $M = \{(g, s_g) : g \in M, s_g \in \{-1, +1\}\}$, module activity is the signed mean of standardized gene values,

$$S_{iM}(t) = \frac{1}{|M|} \sum_{g \in M} s_g \, z_{ig}(t), \tag{S17}$$

where $z_{ig}(t)$ is the standardized projected gene value. Route-level curves average $S_{iM}(t)$ over cells assigned to the relevant source route or branch cohort. Temporal module classes in the main figures are assigned from the shape of these route-mean curves, for example early-peaking, route-retained or late-rising patterns.

For the *C. elegans* terminal-selector validation, we additionally computed a lagged conditional-mutual-information directionality score on generated branch trajectories. Generated endpoints were assigned to ciliated or non-ciliated neuronal terminal branches using the same $k$-nearest-neighbour state-assignment procedure. For each documented regulatory relationship, projected gene values were evaluated along the generated trajectories assigned to the relevant branch. For each delay $d \in \{1, 2, 3\}$, we estimated the forward conditional mutual information

$$I(r_t; q_{t+d} \mid q_{t+d-1}), \tag{S18}$$

where $r$ is the regulator gene score and $q$ is the target gene score, and compared it with the reverse quantity $I(q_t; r_{t+d} \mid r_{t+d-1})$. The reported directionality score is the largest forward value across delays minus the largest reverse value across delays. Positive values indicate stronger lagged conditional information in the known regulator-to-target direction. Activating and repressive relationships are both evaluated by this directional criterion. The local influence and velocity-network analyses below quantify the sensitivity of future gene or module scores to local perturbations of the trained velocity field.

The same directionality score was used for the documented regulatory relationship panels shown in Supplementary Figs. S5 and S10. In *C. elegans*, 30 benchmark relationships were assembled from the neuronal-development literature together with WormBook and WormBase gene-function annotations. The panel covers GABAergic terminal identity, DAF-19/RFX-associated ciliogenesis, the UNC-86–MEC-3 touch-receptor program, UNC-3-associated cholinergic motor-neuron identity, neurogenic priming and late neuronal-selector programs [19–29]. Relationships were evaluated when both genes were represented in the ABpxp trajectory analysis and assigned to the corresponding documented developmental branch.

In the LARRY hematopoiesis analysis, 72 benchmark relationships were assembled from signed TRRUST v2 transcriptional-regulatory entries and literature-supported myeloid regulator-program relationships [30–35]. The TRRUST component comprised signed mouse or human TF-target interactions for hematopoietic regulators. The literature component represented monocyte, inflammatory-myeloid and granulocytic programs present in the LARRY gene set. Relationships were evaluated when both genes were represented in the trajectory analysis. In both datasets, each relationship was scored in its annotated branch or founder context. The recovery fraction is the proportion of relationships with positive directionality in the expected regulator-to-target or regulator-to-program direction.

Local influence and velocity-network panels summarize sensitivities of the trained velocity field. At a representative trajectory state $\mathbf{x}$, a small signed perturbation $\epsilon d_u$ is applied in PCA feature space along an input gene or module direction, mapped through the encoder, and advanced by one short velocity step of length $h$. The local influence from input direction $u$ to output quantity $v$ is

$$J_{v,u}(\mathbf{x}) = \frac{R_v(\theta[\mathbf{z} + h\,\mathbf{v}_\psi(\mathbf{z}_u, t)]) - R_v(\theta[\mathbf{z} + h\,\mathbf{v}_\psi(\mathbf{z}, t)])}{\epsilon}, \tag{S19}$$

where $\mathbf{z} = \phi(\mathbf{x})$, $\mathbf{z}_u = \phi(\mathbf{x} + \epsilon d_u)$ and $R_v$ projects decoded PCA features to the output gene or module. The reported local gene- and module-sensitivity matrices average $J_{v,u}$ over founder, intermediate or late route states; branch-differential edges subtract the matrix for one route from the matched matrix for the other route.

Spatial-neighbourhood analyses in eTracer use tissue coordinates within each section. For each source cell, the eight nearest neighbours define local fractions of macrophage, fibroblast, endothelial, S3 and S4 cells. The S3-state probability is evaluated after integrating the source cell to the next section. Cells are grouped by the upper and lower quartiles of this probability, and neighbourhood enrichment compares mean local cell-state fractions between the two groups. Ridge regression summarizes associations with local cell-state composition, radial position, inferred growth and AP-1 score. Moran's $I$ on the same neighbour graph quantifies spatial autocorrelation of the inferred S3-state probability.

**Algorithm S3** PhyloFM inference: joint latent-and-mass integration

**Require:** Initial population $\mathbf{x}^{(0)}$; trained $(\phi, \theta, \mathbf{v}_\psi, g_\xi)$; evaluation times $0 = t_0 < t_1 < \cdots < t_L = T$
**Ensure:** Latent trajectory $\mathbf{z}(t)$, log-mass $\log m(t)$, decoded PCA feature vector $\hat{\mathbf{x}}(t)$ at evaluation times

$\mathbf{z}(0) \leftarrow \phi(\mathbf{x}^{(0)})$; $\log m(0) \leftarrow 0$
**for** $\ell = 0, 1, \ldots, L-1$ **do**
  $\big(\mathbf{z}(t_{\ell+1}), \log m(t_{\ell+1})\big) \leftarrow \text{IntegratorStep}(\mathbf{v}_\psi, g_\xi, \mathbf{z}(t_\ell), \log m(t_\ell), t_\ell, t_{\ell+1})$
**end for**
$\hat{\mathbf{x}}(t) \leftarrow \theta(\mathbf{z}(t))$ at the requested evaluation times
**return** $\big\{\mathbf{z}(t), \log m(t), \hat{\mathbf{x}}(t)\big\}$

In all analyses reported here, IntegratorStep was implemented with the adaptive Dormand–Prince ODE solver in `torchdiffeq` (`dopri5`), using relative and absolute tolerances of $10^{-5}$ and $10^{-7}$ unless otherwise stated.

#### S3.5 Per-dataset hyperparameters

Tables S1–S5 list the hyperparameter values used for each benchmark. Architecture and learning-rate settings are shared across datasets. Dataset-specific values include loss weights, OT regularization and, for eTracer, spatial representation and coupling parameters. The "Topology source" row identifies the embryonic lineage tree or CARTA-derived differentiation graph used in each analysis.

Table S1: PhyloFM training hyperparameters for the *C. elegans* benchmark.

| Parameter | Value |
|---|---|
| *Architecture* | |
| Encoder / decoder | 5-layer residual MLP |
| Hidden dimension | 256 |
| Latent dimension | 64 |
| Encoder input / decoder target | PCA components |
| Topology source | exact developmental tree |
| Random seed | 0 |
| *Representation (VAE) phase* | |
| Optimizer | Adam |
| Learning rate | $5\times10^{-4}$ |
| Epochs | 16,000 |
| $\lambda_{\text{rec}}$ (reconstruction weight) | 1.0 |
| $\lambda_{\text{top}}$ (lineage weight) | 2.0 |
| KL warm-up | linear, $\beta : 0 \to 0.1$ over 16,000 epochs |
| *Dynamics (flow-matching) phase* | |
| Optimizer | Adam |
| Learning rate | $5\times10^{-3}$ |
| Epochs | 3000 |
| Batch size | 512 |
| $\alpha$ (latent vs. PCA feature mix) | 0.9 |
| OT entropy regularization $\varepsilon$ | 0.05 |
| UOT marginal regularization $\rho$ | 1.0 |
| Blockwise UOT subproblem size | 1,000 |
| Growth weight $\lambda_g$ | 1.0 |

Table S2: PhyloFM training hyperparameters for the LARRY hematopoiesis benchmark.

| Parameter | Value |
|---|---|
| *Architecture* | |
| Encoder / decoder | 5-layer residual MLP |
| Hidden dimension | 256 |
| Latent dimension | 64 |
| Encoder input / decoder target | PCA components |
| Topology source | CARTA-derived hematopoietic differentiation graph |
| Random seed | 0 |
| *Representation (VAE) phase* | |
| Optimizer | Adam |
| Learning rate | $5 \times 10^{-4}$ |
| Epochs | 16,000 |
| $\lambda_{\text{rec}}$ | 1.5 |
| $\lambda_{\text{top}}$ | 2.0 |
| KL warm-up | linear, $\beta : 0 \to 0.1$ over 16,000 epochs |
| *Dynamics (flow-matching) phase* | |
| Optimizer | Adam |
| Learning rate | $5 \times 10^{-3}$ |
| Epochs | 3000 |
| Batch size | 512 |
| $\alpha$ (latent vs. PCA feature mix) | 0.9 |
| OT entropy regularization $\varepsilon$ | 0.2 (T2$\to$T4), 0.1 (T4$\to$T6) |
| UOT marginal regularization $\rho$ | 3.0 |
| Blockwise UOT subproblem size | 1,000 |
| Growth weight $\lambda_g$ | 1.0 |

Table S3: PhyloFM training hyperparameters for the pandaCREST benchmark.

| Parameter | Value |
|---|---|
| *Architecture* | |
| Encoder / decoder | 5-layer residual MLP |
| Hidden dimension | 256 |
| Latent dimension | 64 |
| Encoder input / decoder target | PCA components |
| Topology source | CARTA-derived ventral-midbrain differentiation graph |
| Random seed | 0 |
| *Representation (VAE) phase* | |
| Optimizer | Adam |
| Learning rate | $5\times10^{-4}$ |
| Epochs | 16,000 |
| $\lambda_{\mathrm{rec}}$ | 1.5 |
| $\lambda_{\mathrm{top}}$ | 2.0 |
| KL warm-up | linear, $\beta : 0 \to 0.1$ over 16,000 epochs |
| *Dynamics (flow-matching) phase* | |
| Optimizer | Adam |
| Learning rate | $5\times10^{-3}$ |
| Epochs | 3000 |
| Batch size | 512 |
| $\alpha$ (latent vs. PCA feature mix) | 0.9 |
| OT entropy regularization $\varepsilon$ | 0.1 |
| UOT marginal regularization $\rho$ | 1.0 |
| Blockwise UOT subproblem size | 1,000 |
| Growth weight $\lambda_g$ | 1.0 |

Table S4: PhyloFM training hyperparameters for the zebrafish heart regeneration benchmark.

| Parameter | Value |
|---|---|
| *Architecture* | |
| Encoder / decoder | 5-layer residual MLP |
| Hidden dimension | 256 |
| Latent dimension | 64 |
| Encoder input / decoder target | PCA components |
| Topology source | CARTA-derived 14-state cardiac differentiation graph |
| Random seed | 0 |
| *Representation (VAE) phase* | |
| Optimizer | Adam |
| Learning rate | $5\times10^{-4}$ |
| Epochs | 16,000 |
| $\lambda_{\text{rec}}$ | 5.0 |
| $\lambda_{\text{top}}$ | 2.0 |
| KL warm-up | linear, $\beta : 0 \to 0.1$ over 16,000 epochs |
| *Dynamics (flow-matching) phase* | |
| Optimizer | Adam |
| Learning rate | $5\times10^{-3}$ |
| Epochs | 3000 |
| Batch size | 512 |
| $\alpha$ (latent vs. PCA feature mix) | 0.9 |
| OT entropy regularization $\varepsilon$ | 0.05 |
| UOT marginal regularization $\rho$ | 1.0 |
| Blockwise UOT subproblem size | 1,000 |
| Growth weight $\lambda_g$ | 1.0 |

Table S5: PhyloFM training hyperparameters for the eTracer spatiotemporal benchmark.

| Parameter | Value |
|---|---|
| *Architecture* | |
| Encoder / decoder | 5-layer residual MLP |
| Hidden dimension | 256 |
| Latent dimension | 64 |
| Encoder input / decoder target | PCA components |
| Topology source | CARTA-derived tumour-state graph with spatial neighbourhoods |
| Random seed | 0 |
| *Representation (VAE) phase* | |
| Optimizer | Adam |
| Learning rate | $5\times10^{-4}$ |
| Epochs | 16,000 |
| $\lambda_{\text{rec}}$ | 1.5 |
| $\lambda_{\text{top}}$ | 2.0 |
| KL warm-up | linear, $\beta : 0 \to 0.1$ over 16,000 epochs |
| *Dynamics (flow-matching) phase* | |
| Optimizer | Adam |
| Learning rate | $5\times10^{-3}$ |
| Epochs | 3000 |
| Batch size | 1024 |
| $\alpha$ (latent vs. PCA feature mix) | 0.9 |
| OT entropy regularization $\varepsilon$ | 0.05 |
| UOT marginal regularization $\rho$ | 1.0 |
| Blockwise UOT subproblem size | 4,000 |
| Growth weight $\lambda_g$ | 1.0 |
| *Spatial terms (eTracer only)* | |
| Within-slice spatial neighbours $k$ | 8 |
| Within-slice spatial weight $\lambda_{\text{sp}}$ | 0.1 |
| Feature term weight (PCA + latent) | 0.7 |
| Raw spatial-coordinate term weight | 0.3 |

### Supplementary Note S4. Benchmark design, baselines and metrics

This note describes the trajectory-inference methods compared with PhyloFM in the main text and summarizes how each method represents cell-state change and population size. The four non-spatial baselines use time-resolved molecular profiles. The eTracer benchmarks additionally incorporate tissue coordinates.

#### S4.1 Non-spatial baselines

**OT-CFM** [15] pairs cells from adjacent snapshots by minibatch optimal transport in the input feature space. The paired cells define conditional paths used to train a deterministic velocity field.

**VGFM** [12] couples adjacent snapshots with semi-relaxed optimal transport and interprets the resulting plan as population-size adjustment followed by cell-state movement. Separate neural networks learn velocity and growth from conditional matching targets derived from the coupling. A Wasserstein distribution loss on integrated trajectories further aligns the generated and observed populations.

**TIGON** [36] describes population dynamics with a velocity field and a growth field under the Wasserstein–Fisher–Rao formulation of dynamical optimal transport. Separate neural networks represent these fields and are trained through neural ODE integration so that the generated populations match the observed snapshots. We applied TIGON to the same PCA features used for the other dynamical baselines.

**scDiffEq** [37] represents cell-state dynamics with a neural stochastic differential equation. Neural networks learn the deterministic drift and state-dependent diffusion. Training advances cells from the earliest snapshot and minimizes the Sinkhorn divergence between generated and observed populations at later times.

#### S4.2 Spatial baselines

**STORIES** [38] learns a scalar differentiation potential from spatial transcriptomic time series. The gradient of this potential defines cell-state movement, and a fused Gromov–Wasserstein loss compares molecular profiles together with spatial organization. The learned potential generates trajectories and a spatially informed differentiation index. Population weights can be derived from proliferation and apoptosis gene scores; uniform weights were used in the eTracer benchmark.

**stVCR** [39] uses dynamical unbalanced optimal transport to reconstruct molecular differentiation, population growth and movement in physical space. It also estimates rigid transformations between spatial coordinate systems when sections are not aligned. The model learns separate molecular velocity, spatial velocity and growth fields from time-resolved spatial transcriptomic data.

#### S4.3 Benchmark implementation

All baseline models were run with author-provided implementations and author-recommended settings on the same processed expression representation.

#### S4.4 Benchmark metrics

All benchmark metrics were computed after converting each method's output to the same 50-dimensional PCA feature space used by PhyloFM. For a source population integrated to evaluation time $t_k$, let $\hat{\mathbf{x}}_i(t_k)$ be the predicted PCA state and $\hat{w}_i(t_k)$ its predicted mass. The observed target population is represented by PCA states $\mathbf{x}_j^{(k)}$ with uniform weights. Population transport error was measured by exact optimal transport between these two empirical measures. W1 used the Euclidean cost matrix $C_{ij} = \|\hat{\mathbf{x}}_i(t_k) - \mathbf{x}_j^{(k)}\|_2$,

$$\mathrm{W1}(t_k) = \min_{\gamma \in \Pi(\hat{a}, b)} \sum_{ij} \gamma_{ij} C_{ij}, \tag{S20}$$

where $\hat{a}$ is the normalized predicted mass vector and $b$ is uniform over the observed target cells. W2 used the squared-Euclidean cost and reported the square root of the optimal value,

$$\mathrm{W2}(t_k) = \left[ \min_{\gamma \in \Pi(\hat{a}, b)} \sum_{ij} \gamma_{ij} \|\hat{\mathbf{x}}_i(t_k) - \mathbf{x}_j^{(k)}\|_2^2 \right]^{1/2}. \tag{S21}$$

Methods that return state trajectories alone were evaluated with uniform source-cell weights. In supporting benchmark tables we also report total-mass variation, defined as $|\sum_i \hat{w}_i(t_k) - n_k/n_0|/(n_k/n_0)$, where $n_0$ and $n_k$ are the observed source and target cell counts.

Non-holdout transport scores used models trained on all adjacent sampled intervals. PhyloFM, VGFM, TIGON and scDiffEq were evaluated by integrating cells from the first sampled time point to each target time. The reported OT-CFM transport scores use rollouts spanning at most two consecutive intervals. For the *C. elegans* time-point holdout benchmark, each interior stage (210, 270, 330, 390 and 450 min) was omitted in turn. Each method was retrained for every split. In PhyloFM, cells from the omitted stage were excluded from representation learning and flow matching, while the common PCA coordinate system was retained across splits. A bridge coupling between the flanking observed stages allowed the learned field to cross the missing interval. Predictions at the omitted stage were scored by W1 and W2, and the reported values are the mean across the five splits. The lineage-local dynamic error described below was computed in the non-holdout adjacent-time analysis.

The lineage-local dynamic error in the *C. elegans* benchmark uses the known embryonic tree to form a ground-truth coupling $Q^{(k)}$ between adjacent stages. For each source cell, $Q_{ij}^{(k)}$ gives the normalized distribution of its true descendants or descendant-compatible target cells at the next sampled time. The local one-step error is

$$\mathrm{DynErr}(k) = \sum_i \omega_i \sum_j Q_{ij}^{(k)} \left\| \hat{\mathbf{x}}_i(t_{k+1}) - \mathbf{x}_j^{(k+1)} \right\|_2^2, \tag{S22}$$

with source weights $\omega_i$ normalized over rows that have nonzero ground-truth support. The reported value is the mean over adjacent time gaps.

Clone-grounded fate metrics use the fate-probability procedure defined in Supplementary Note S3.4. For each clone $c$, the predicted fate distribution is the mean of the predicted per-cell distributions over source cells in that clone,

$$\hat{F}_c(r) = \frac{1}{|S_c|} \sum_{i \in S_c} p_i(r \mid t), \tag{S23}$$

and the observed distribution $F_c(r)$ is the fraction of target descendants from the same clone assigned to fate $r$. In LARRY, the primary endpoint score is top-1 fate accuracy across 10 fate classes, with monocyte-versus-neutrophil subsets retained as supporting analyses. In pandaCREST, same-state hidden-future accuracy is the Spearman and Pearson correlation between the predicted $\mathrm{GLU^{BM}}$-minus-$\mathrm{GABA^{BL}}$ fate margin and the empirical clone-derived $\mathrm{GLU^{BM}}$-minus-$\mathrm{GABA^{BL}}$ margin on

NPBM source cells. In eTracer, clone-grounded reconstruction concordance compares inferred and observed distributions over S2, S3 and S4 tumour states. Dominant-state agreement is accompanied by cross-entropy, Brier score, per-state Spearman correlation and mean absolute error.

For the LARRY endpoint benchmark shown in Supplementary Fig. S7, day-2 source cells were propagated to day 6 in the common PCA feature space and assigned a terminal fate by $k$-nearest-neighbour voting against observed day-6 cells ($k = 20$). The assigned fates were compared with clone-derived labels for 859 day-2 source cells spanning all annotated terminal fates.

#### Supplementary Note S5. Lineage-record formats and anchor-topology rationale

The five datasets span the main formats in which ancestry is currently recorded. These include an exact developmental tree in *C. elegans* [40, 41], shared lentiviral barcodes in LARRY hematopoiesis [42], clone-split CRISPR recorders in pandaCREST [43], CRISPR scars recorded independently in each zebrafish [44, 45] and spatially resolved clone labels in eTracer [46]. Each format supports a different set of comparisons across time and biological samples, as summarized in Table S6.

PhyloFM represents each lineage format as a topology over cell-state anchors before learning dynamics. Shared barcodes contribute clone relationships within the experiments in which they were generated. Sample-specific scars and evolving edits contribute through the cell-state graph assembled from their lineage structure. The resulting topology provides a common input for learning velocity, growth and fate probabilities.

Table S6: Lineage records used in this study and the comparisons each one supports. "Across time" indicates whether the same lineage label, when found at two sampled times in the experiment, identifies the same clonal lineage. "Spatial" indicates whether tissue coordinates are recorded jointly with the lineage label. "Evaluation signal" lists the lineage-based comparison used in the study. Within-animal lineage trees, anchor-graph construction and the eTracer static-vs-evolving distinction are discussed in the per-dataset paragraphs below.

| **Dataset** | **Lineage record** | **Across time** | **Spatial** | **Evaluation signal** |
|---|---|---|---|---|
| *C. elegans* ABpxp [40, 41] | invariant embryonic lineage tree from microscopy | yes | no | lineage-local dynamic error |
| LARRY hematopoiesis [42] | shared lentiviral barcodes, in vitro | yes | no | clone-derived mono vs. neutrophil bias at day-2 founders |
| pandaCREST mouse vMB [43] | dual-recorder (V1+V2) CRISPR barcodes; clone-split E11.5 → day-7 organoid | yes | no | clone-derived $\text{GLU}^{\text{BM}}$ vs. $\text{GABA}^{\text{BL}}$ fate margin (NPBM cells) |
| Zebrafish heart regen. (LINNAEUS) [44, 45] | CRISPR scars in zebrabow dTomato, fixed by gastrulation | no | no | fibroblast-branch identity at 0/3/7 dpi |
| eTracer LUAD [46] | static SB1 and SB2 clone barcodes *plus* evolving 3′UTR edits, with Stereo-seq expression profiles | yes (static); no (evolving) | yes | clone-grounded S2-to-S2/S3/S4 state concordance |

**Exact embryonic tree (*C. elegans*).** The Sulston tree assigns every sampled cell to a defined position in an animal-invariant lineage [40]. Combined with the lineage-resolved *C. elegans* embryogenesis atlas [41], this format supports comparison of lineage identity across all sampled times and between animals. We use the tree directly as the anchor graph. The lineage-local dynamic error reported in the main text evaluates predictions using both developmental time and ancestor identity.

**Shared barcodes in vitro (LARRY).** In LARRY, lentiviral integration tags a hematopoietic founder cell, and its descendants inherit the same barcode [42]. Aliquots from the same labelled culture are profiled at several days, so a barcode observed at day 2 and again at day 4 or day 6 identifies the same clone over time. This provides clone-level fate information for the founder population. Barcode identities are specific to the labelling experiment, and comparisons beyond that experiment are made through shared cell-state anchors.

**Clone-split CRISPR recorders (pandaCREST).** pandaCREST uses heritable CREST V1+V2 barcodes whose recording becomes stable approximately 12 h after Cre-driven Cas9 activation in vivo [43]. Half of the E11.5 ventral-midbrain cells from each embryo are profiled directly. The remaining cells are cultured for one week as a brain organoid and then profiled. Shared barcodes connect the two sister populations created by this split and provide the day-7 fate composition associated with each E11.5 clone. Because each embryo generates its own barcode set, clone comparisons are made within embryos. PhyloFM uses the resulting clone-level futures to evaluate fate bias among E11.5 cells with similar molecular states.

**CRISPR scars recorded separately in each fish (LINNAEUS-style zebrafish).** In the Hu et al. zebrafish heart regeneration dataset [45], Cas9 and an sgRNA targeting the multi-copy dTomato locus of the zebrabow line are injected into one-cell-stage embryos. Scar barcodes accumulate during early development and are subsequently retained throughout the life of the animal [44]. Each adult fish carries a distinct scar set, and the 0, 3 and 7 dpi samples were collected from different individuals. Within each fish, the scars define a tree connecting the jointly sampled cells. CARTA combines these per-fish trees into a shared graph over cell-state anchors, which provides the topology used for cross-time dynamical inference.

**Spatial clone records (eTracer).** eTracer combines a CRISPR-Cas9 lineage tracer with Stereo-seq spatial transcriptomics [46]. Static SB1 and SB2 barcodes are introduced into the parental EP-eTracer population and inherited by tumour cells. A second component uses constitutive and Dox-inducible sgRNAs to generate evolving edits in highly expressed endogenous genes as cells divide. Stereo-seq captures the transcriptome, both lineage components and tissue coordinates from the same library. The static barcodes preserve clone identity across the BL, EE and LE tumours of one experimental series and provide clone-level state composition for the reconstruction-concordance analysis. The evolving edits accumulate after engraftment and describe lineage structure within each tumour. PhyloFM incorporates this within-tumour structure through a CARTA-derived anchor graph. Tissue coordinates additionally constrain local latent geometry and contribute to the spatial term in the hybrid coupling.

#### Supplementary Note S6. Capability matrix of PhyloFM and related methods

Table S7 summarizes the biological inputs and dynamical outputs of lineage-structured methods, continuous dynamical models and spatial optimal-transport frameworks.

Table S7: Capability matrix of PhyloFM and related methods. "Lineage prior" indicates whether lineage information contributes to model inference. "Continuous field" indicates whether the model learns a continuous dynamical field that can be integrated between sampled times. "Growth field" indicates whether the model learns a state-dependent population-mass field. "Spatial coordinates" indicates whether measured tissue locations can enter the model.

| **Method** | **Lineage prior** | **Continuous field** | **Growth field** | **Spatial coordinates** | **Principal inferred object** |
|---|---|---|---|---|---|
| **PhyloFM (this work)** | yes | yes | yes | yes | topology-shaped latent geometry, population coupling, fate probabilities, velocity and growth fields |
| LineageOT [47] | yes | no | no | no | ancestry-informed coupling between sampled times |
| Moslin [48] | yes | no | no | no | cross-time coupling from molecular and lineage distances |
| moscot [49] | no | no | no | optional | optimal-transport mapping across time, conditions or space |
| CoSpar [50] | yes | no | no | no | clone-informed fate-transition map |
| CARTA [51] | yes | no | no | no | differentiation graph over potency-defined progenitor and observed terminal cell types |
| OT-CFM [15] | no | yes | no | no | flow-matched velocity field |
| VGFM [12] | no | yes | yes | no | velocity and growth fields |
| TIGON [36] | no | yes | yes | no | velocity and growth fields |
| scDiffEq [37] | no | yes | no | no | drift and diffusion fields |
| STORIES [38] | no | yes | no | yes | spatially informed differentiation potential |
| stVCR [39] | no | yes | yes | yes | molecular velocity, spatial velocity and growth fields |

PhyloFM separates topology construction from dynamical inference. The known embryonic tree or a CARTA-derived differentiation graph defines distances between cell-state anchors. Representation learning embeds these distances into a continuous geometry used to estimate adjacent-time couplings and learn velocity and growth fields.

Several methods address related lineage questions. LineageVAE reconstructs unobserved progenitor states and historical transcriptomes from cells with shared barcodes [52]. Deep Lineage combines an autoencoder with recurrent networks to predict expression at unobserved times and early fate bias within clones [53]. Clone2vec represents variation among clones and its association with gene expression [54]. These approaches provide complementary views of historical cell states, clone-conditioned prediction and clonal organization.

Continuous benchmarks use OT-CFM, VGFM, TIGON and scDiffEq, with STORIES and stVCR used for eTracer. The *C. elegans* analysis also includes static Moslin and LineageOT couplings and flow matching trained from Moslin couplings.

### Supplementary Figure S1. Benchmark robustness and lineage-ablation analysis in *C. elegans*

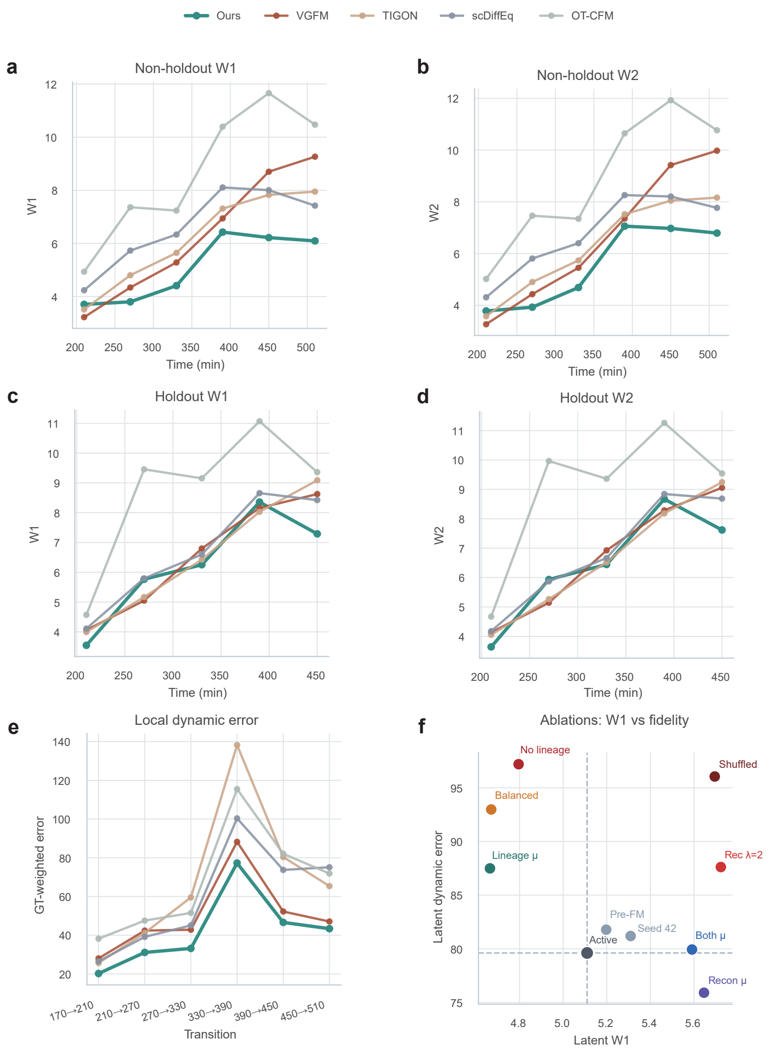


**Supplementary Figure S1.** Benchmark robustness and lineage-ablation analysis in *C. elegans*. **a,b,** Non-holdout W1 and W2 distances across developmental target times for PhyloFM, VGFM, TIGON, scDiffEq and OT-CFM. **c,d,** Time-point holdout W1 and W2 distances after each interior developmental stage was omitted in turn. **e,** Local dynamic error for adjacent exact-lineage transitions from 170 to 510 min. The metric compares model rollouts with the observed next-time population connected by exact lineage. **f,** Ablation analysis comparing latent W1 and latent dynamic error for the reported model and variants that alter lineage, reconstruction and balancing terms. Dashed reference lines mark the reported model. Lower values on both axes indicate better transport fidelity and lineage-local dynamics.

### Supplementary Figure S2. Lineage-aware Moslin-FM and static-coupling comparisons in *C. elegans*

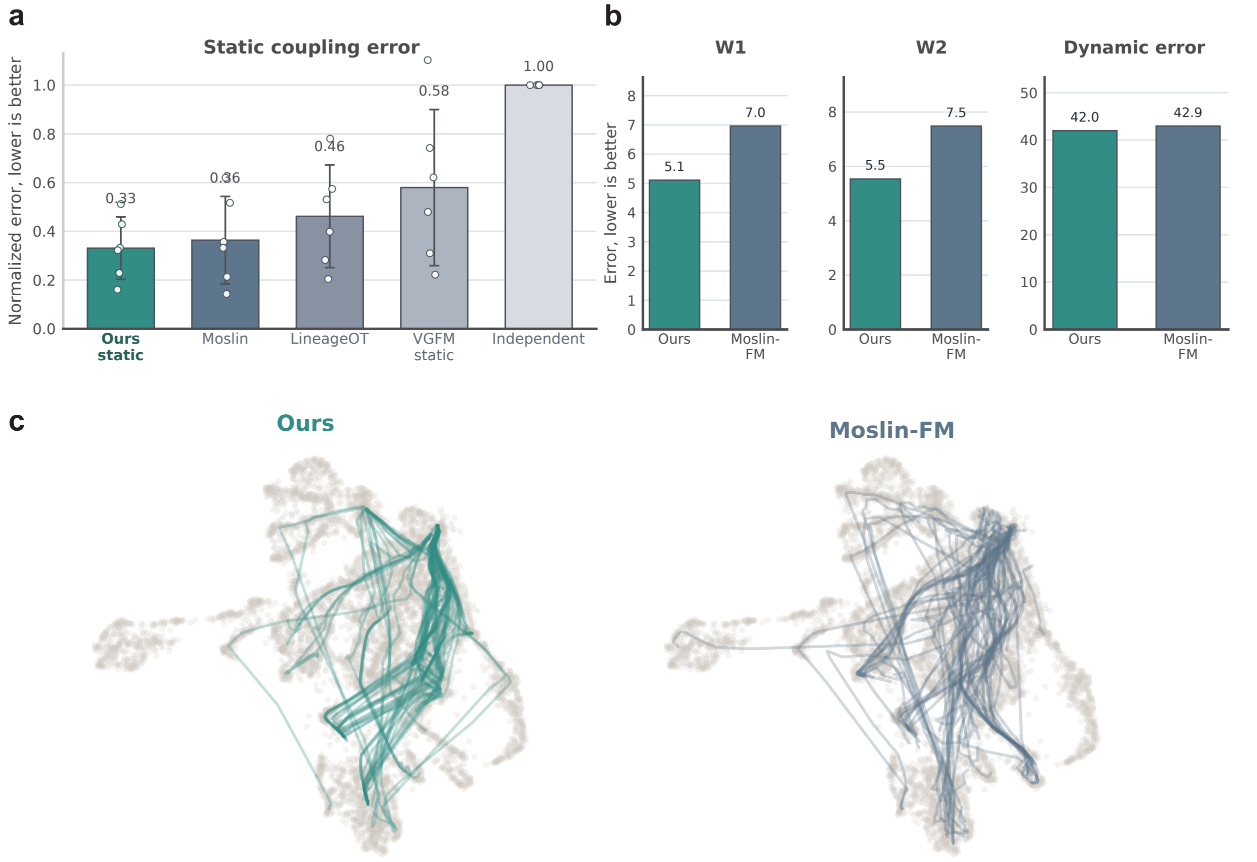


**Supplementary Figure S2.** Lineage-aware Moslin-FM and static-coupling comparisons in *C. elegans*. **a,** Moslin-style normalized static-coupling error for the static training coupling used by PhyloFM, Moslin, LineageOT, the VGFM PCA-flow control and an independent-coupling control. Bars show the mean over adjacent developmental intervals and points show individual intervals. **b,** Final dynamical-model comparison between PhyloFM and Moslin-FM using W1, W2 and lineage-local dynamic error. **c,** UMAP-projected generated trajectories from PhyloFM and Moslin-FM on the observed *C. elegans* manifold. Lower values are better in panels a and b.

#### Supplementary Figure S3. Inferred fine cell-state flow across *C. elegans* neurodevelopment

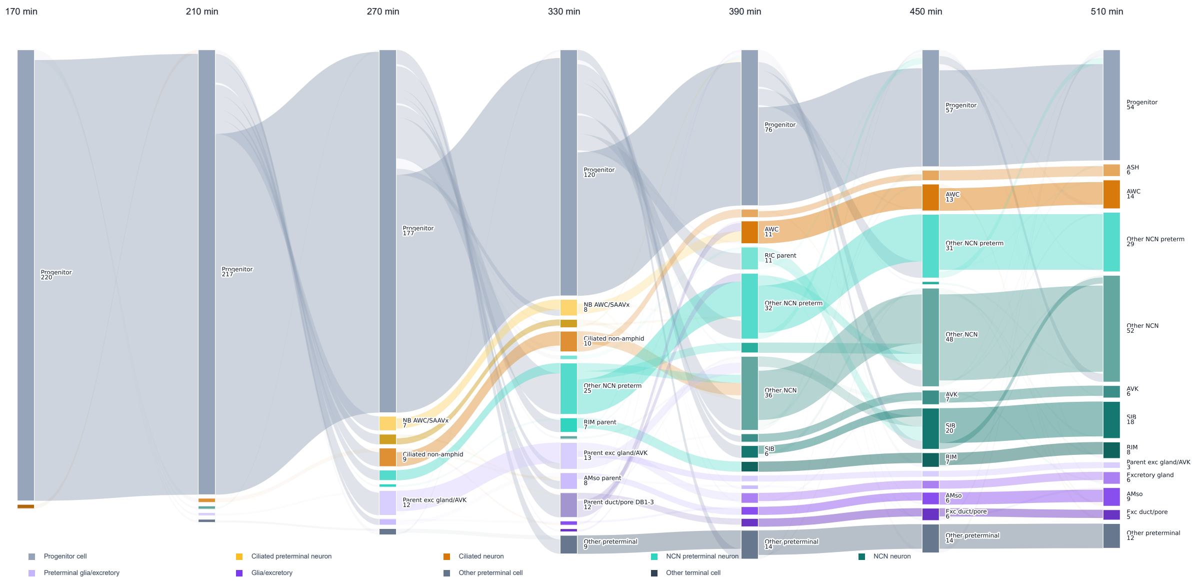


**Supplementary Figure S3.** Inferred fine cell-state flow across *C. elegans* neurodevelopment. Sankey representation of model-inferred trajectories from 170 to 510 min. Each ribbon follows one inferred trajectory. The 170 min nodes are observed states, and later nodes are assigned by hard top-1 nearest-neighbour calls to observed fine cell-state labels. Node heights and labels report trajectory counts at each time point. Colors separate progenitor, ciliated preterminal, ciliated neuron, non-ciliated neuronal preterminal, non-ciliated neuron, glia and excretory, preterminal glia and excretory, other preterminal and other terminal groups. The figure traces progression from an early progenitor-dominated population into AWC and ASH ciliated neurons, non-ciliated neuronal states, glia and excretory classes and remaining preterminal or terminal states.

### Supplementary Figure S4. Module ontology support, founder marker maps and route-specific molecular dynamics

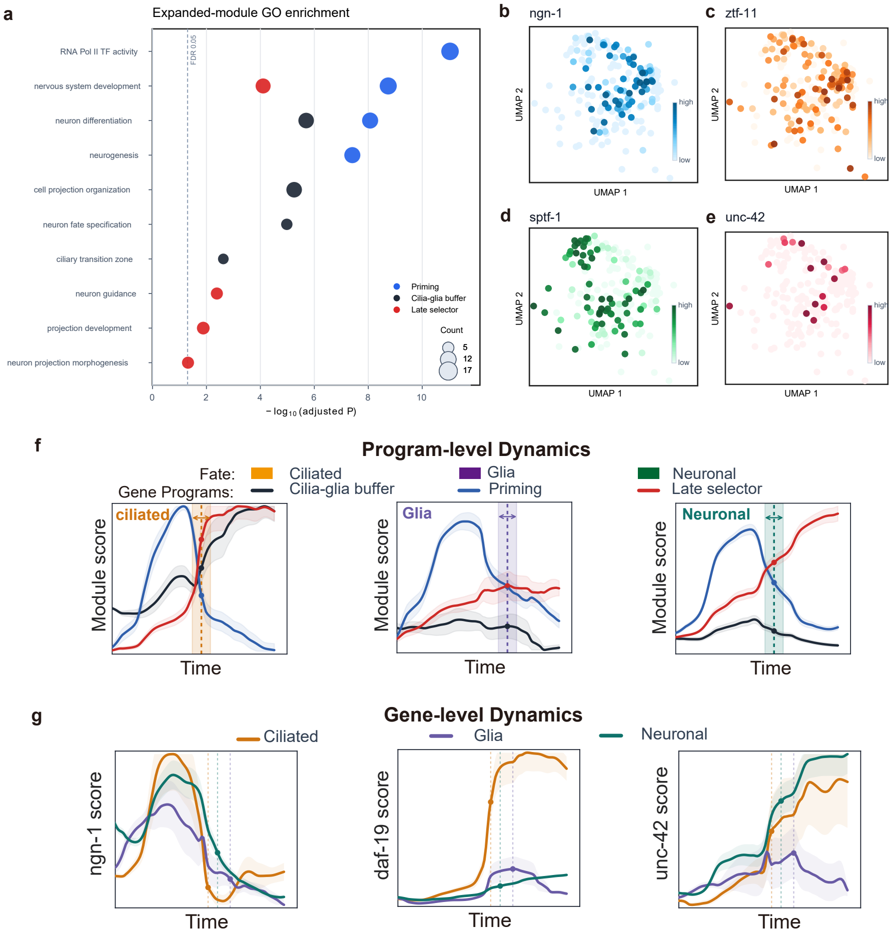


**Supplementary Figure S4.** Module ontology support, founder marker maps and route-specific molecular dynamics. **a,** Gene Ontology enrichment of expanded dynamic-pattern modules. Dot position gives minus log10 adjusted P value, dot size gives the number of genes in the term and color indicates the associated module. The dashed line marks FDR 0.05. **b-e,** Founder-cell UMAP maps colored by marker-gene scores for *ngn-1*, *ztf-11*, *sptf-1* and *unc-42*. Localized score patterns link the priming, cilia-glia buffer and late-selector modules to early founder-state heterogeneity. **f,** Program-level dynamics of the cilia-glia buffer, priming and late-selector modules along ciliated, glial and neuronal trajectories. The priming program rises early and subsequently declines, while the cilia-glia buffer and late-selector programs become more prominent along their associated routes. **g,** Route-specific dynamics of *ngn-1*, *daf-19* and *unc-42*. *ngn-1* peaks early and then declines, *daf-19* rises selectively along the ciliated route, and *unc-42* turns on late and is strongest along the neuronal route. In **f** and **g**, curves show route means and shaded bands show s.e.m.; points and vertical dashed lines mark inferred commitment positions.

### Supplementary Figure S5. Documented regulatory relationships in *C. elegans* neuronal trajectories

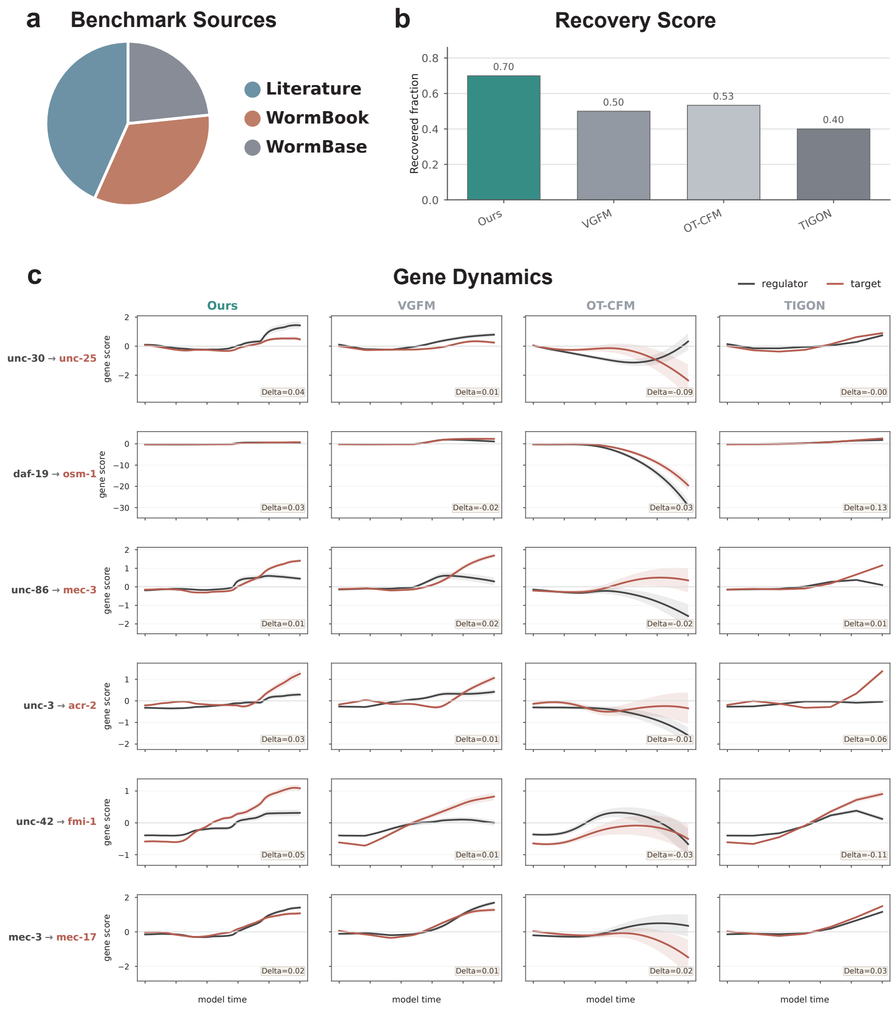


**Supplementary Figure S5.** Documented regulatory relationships in *C. elegans* neuronal trajectories. **a,** Evidence sources for the *C. elegans* regulatory relationship panel. Relationships combine literature, WormBook and WormBase annotations of neuronal terminal programs and were restricted to regulators and targets represented in the analysed ABpxp trajectories. **b,** Recovery fraction under the shared conditional-mutual-information directionality score. A relationship is counted as recovered when the expected regulator-to-target or regulator-to-program direction has a positive directionality score. **c,** Representative dynamics for biologically interpretable examples, including UNC-30-associated GABAergic identity, DAF-19-associated ciliogenesis, UNC-86 and MEC-3-associated touch-receptor specification, UNC-3-associated cholinergic motor identity and UNC-42-associated late neuronal programs. Curves show branch-mean projected gene scores over model time with s.e.m.; black denotes the regulator and red denotes the target or downstream program marker.

#### Supplementary Figure S6. EPIC GFP reporter validation of inferred *C. elegans* gene dynamics

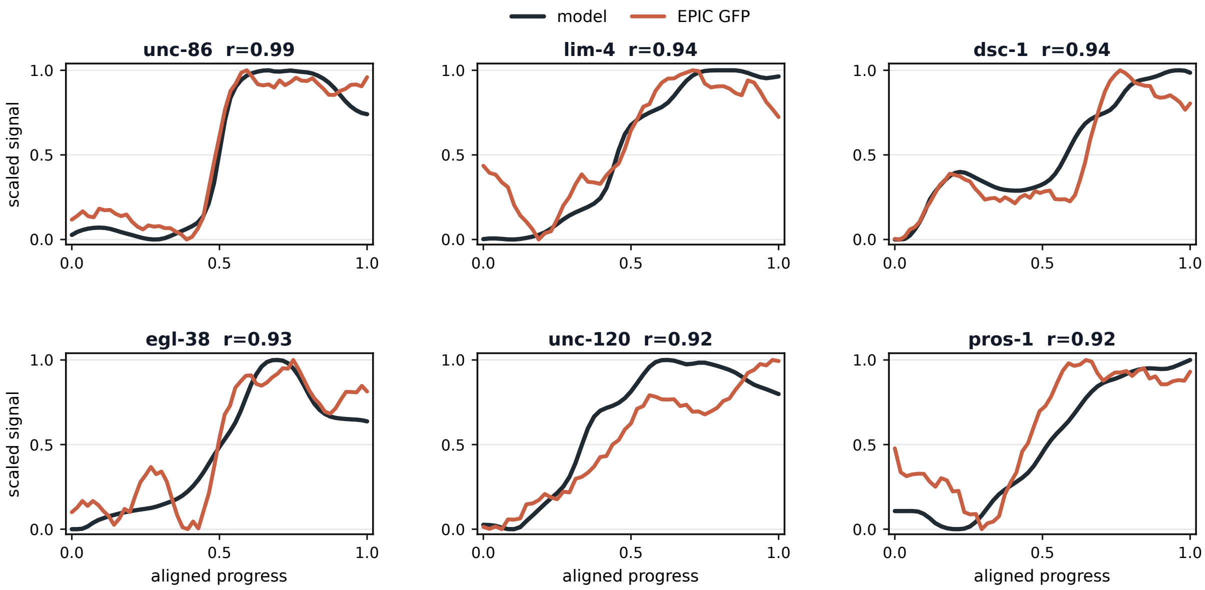


**Supplementary Figure S6.** EPIC GFP reporter validation of inferred *C. elegans* gene dynamics. Six additional reporter examples with high agreement and clear dynamic structure. Black curves show inferred gene dynamics and red curves show matched EPIC GFP reporter traces from the live-embryo expression resource of Murray et al. [55].

#### Supplementary Figure S7. Comparative founder fate maps and accuracy in LARRY hematopoiesis

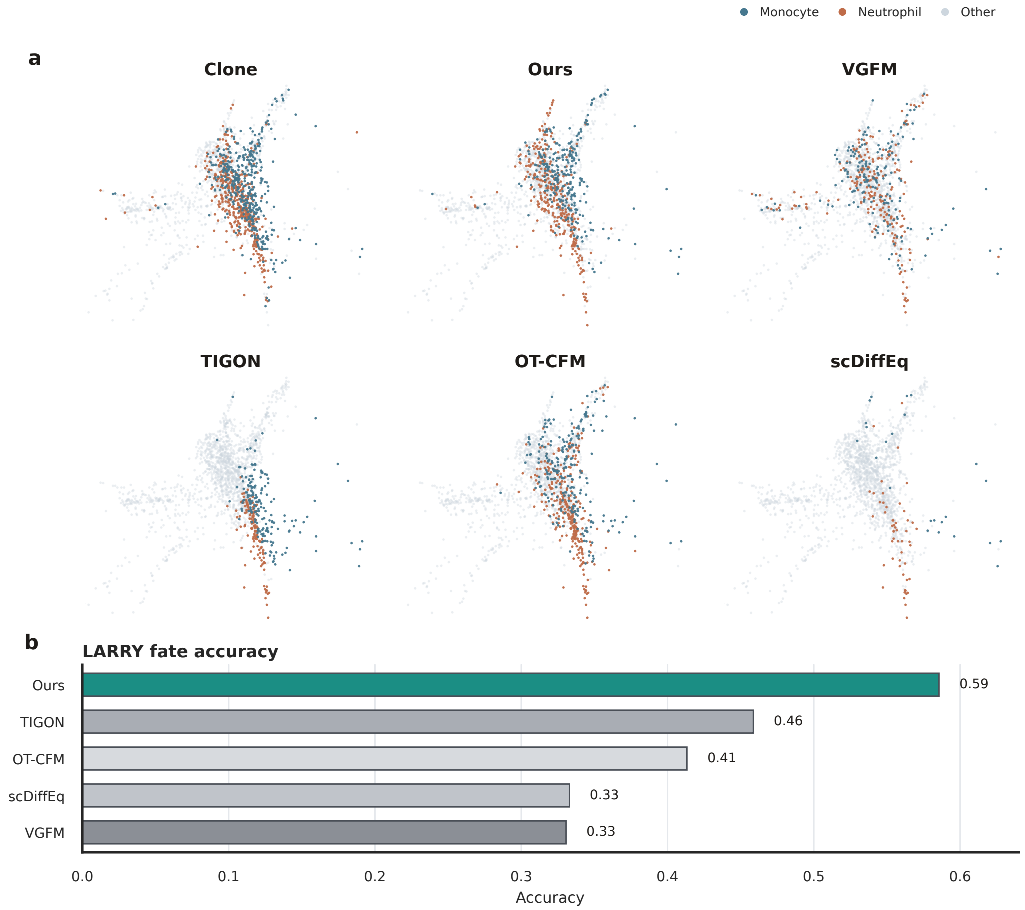


**Supplementary Figure S7.** Comparative founder fate maps and accuracy in LARRY hematopoiesis. **a,** Clone-derived monocyte, neutrophil and other founder labels are shown together with predictions from the model labelled Ours and from VGFM, TIGON, OT-CFM and scDiffEq. **b,** LARRY fate accuracy under the shared terminal-state assignment. The comparison evaluates how well each method recovers future fate structure within the day-2 founder population using shared LARRY barcodes as the reference.

### Supplementary Figure S8. Growth-associated structure in LARRY hematopoiesis

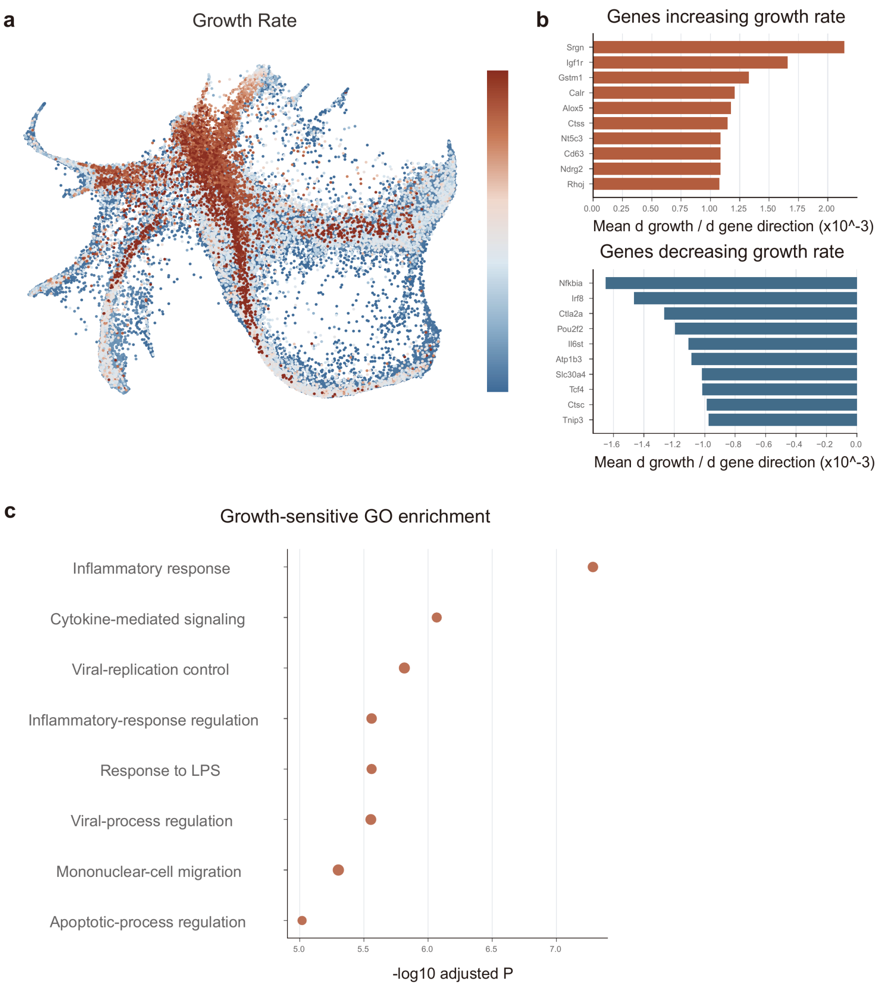


**Supplementary Figure S8.** Growth-associated structure in LARRY hematopoiesis. **a,** Manifold map of inferred growth rate across the hematopoietic landscape. **b,** Genes whose local perturbation directions increase or decrease inferred growth output. *Srgn*, *Igf1r*, *Gstm1*, *Calr*, *Alox5*, *Ctss*, *Nt5c3*, *Cd63*, *Ndrg2* and *Rhoj* increase growth output, while *Nfkbia*, *Irf8*, *Ctla2a*, *Pou2f2*, *Il6st*, *Atp1b3*, *Slc30a4*, *Tcf4*, *Ctsc* and *Tnip3* decrease growth output. **c,** Gene Ontology enrichment for growth-sensitive genes, highlighting inflammatory response, cytokine-mediated signaling, viral-replication control, inflammatory-response regulation, response to LPS, viral-process regulation, mononuclear-cell migration and apoptotic-process regulation.

#### Supplementary Figure S9. Molecular support for hidden founder fate and branch dynamics

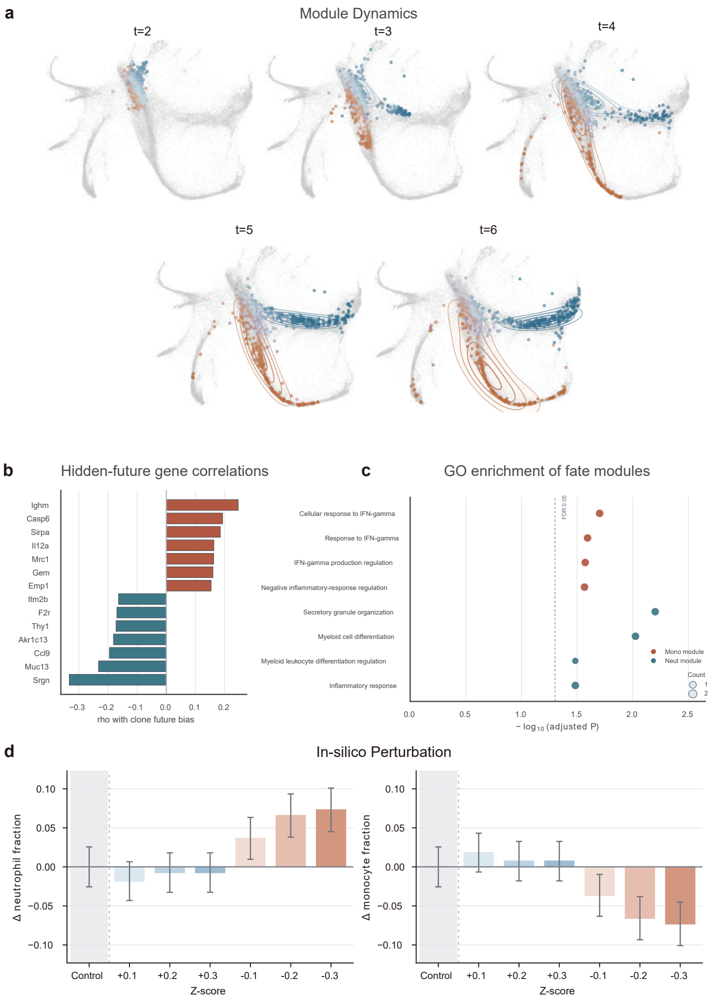


**Supplementary Figure S9.** Molecular support for hidden founder fate and branch dynamics. **a,** Module dynamics from generated time 2 to 6, showing the temporal separation of monocyte- and neutrophil-associated trajectories on the hematopoietic manifold. **b,** Genes correlated with clone-derived future bias. Positive correlations mark genes associated with monocyte-biased futures, and negative correlations mark genes associated with neutrophil-biased futures. **c,** Gene Ontology enrichment of monocyte and neutrophil fate modules. Dot position gives minus log10 adjusted P value, dot size gives the number of genes in the term and color indicates the associated module. The dashed line marks FDR 0.05. **d,** In-silico perturbation of module and regulator scores in mono-biased and neut-biased founders. Bars show changes in terminal neutrophil and monocyte fractions relative to the control condition.

### Supplementary Figure S10. Documented myeloid regulatory relationships in LARRY hematopoiesis

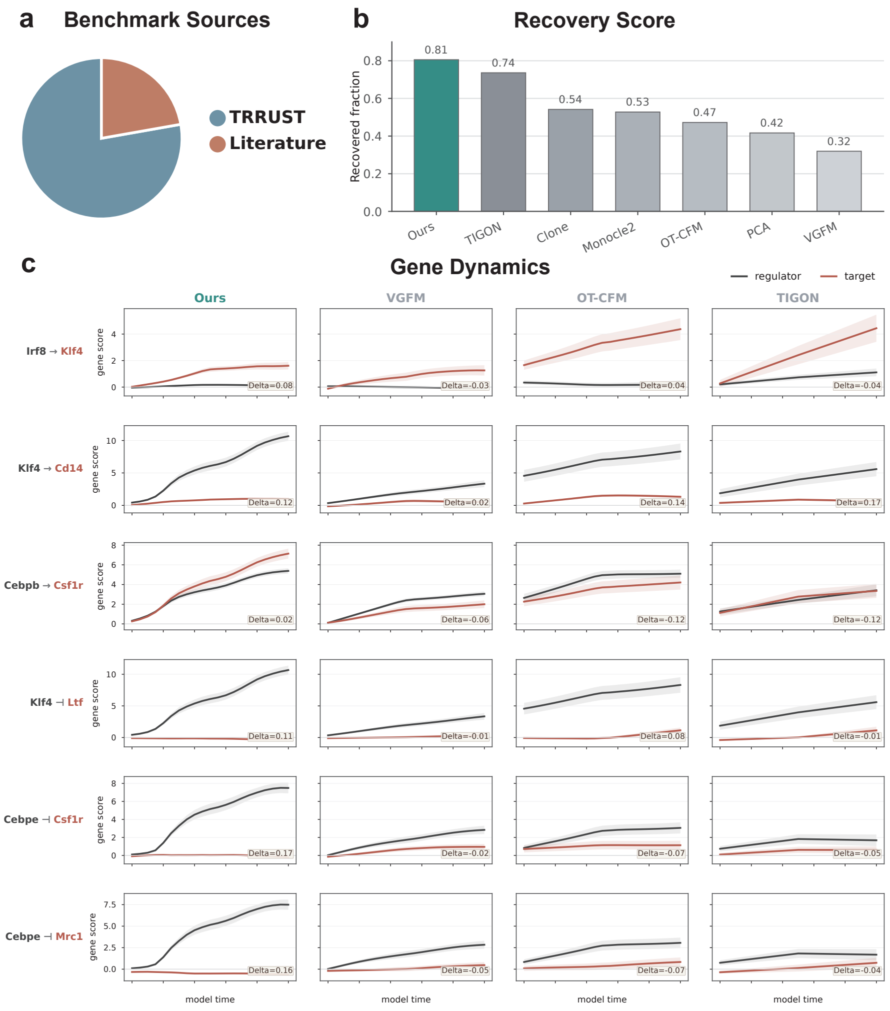


**Supplementary Figure S10.** Documented myeloid regulatory relationships in LARRY hematopoiesis. **a,** Evidence sources for the hematopoietic regulatory relationship panel. Relationships combine TRRUST entries with literature-supported myeloid regulatory programs and were restricted to genes represented in the analysed LARRY trajectories. **b,** Recovery fraction under the shared conditional-mutual-information directionality score. A relationship is counted as recovered when the expected regulator-to-target or regulator-to-program direction has a positive directionality score. **c,** Representative dynamics for myeloid regulatory examples across PhyloFM and baseline models. Curves show branch-mean projected gene scores over model time with s.e.m.; black denotes the regulator and red denotes the target or downstream program marker.

### Supplementary Figure S11. Branch-driver support for pandaCREST NPBM fate structure

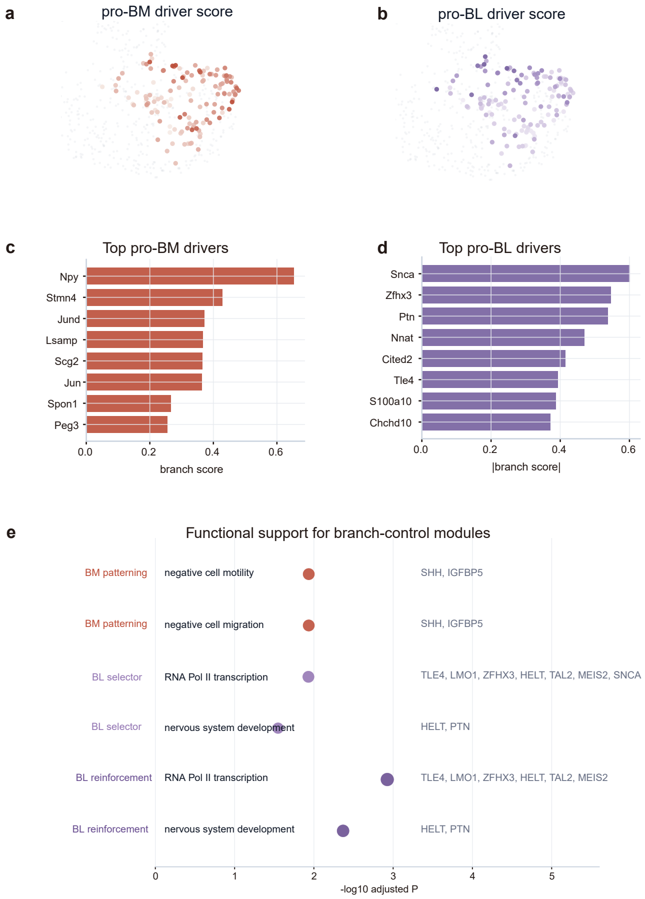


**Supplementary Figure S11.** Branch-driver support for pandaCREST NPBM fate structure. **a,b,** UMAP maps of pro-BM and pro-BL driver scores on NPBM founder cells. **c,d,** Top pro-BM and pro-BL driver genes ranked by branch score. Pro-BM drivers include *Npy*, *Stmn4*, *Jund*, *Lsamp*, *Scg2*, *Jun*, *Spon1* and *Peg3*. Pro-BL drivers include *Snca*, *Zfhx3*, *Ptn*, *Nnat*, *Cited2*, *Tle4*, *S100a10* and *Chchd10*. **e,** Gene Ontology support for branch-control modules. BM-patterning genes are enriched for negative cell motility and migration terms, while BL-selector and BL-reinforcement genes are linked to RNA polymerase II transcription and nervous-system development terms.

### Supplementary Figure S12. Three-module support for the NPBM branch-control model

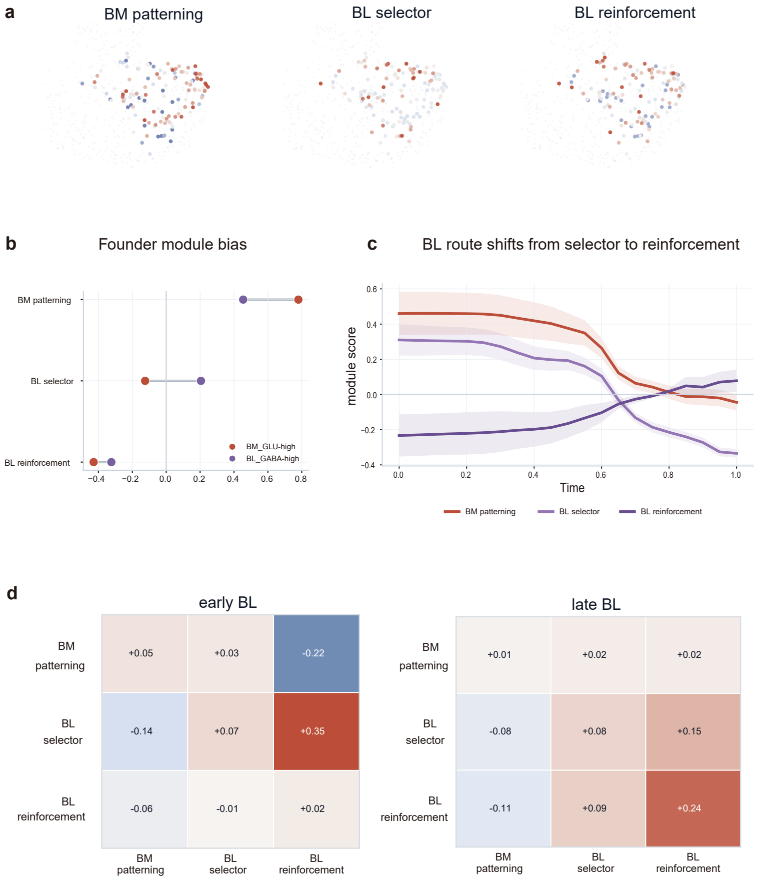


**Supplementary Figure S12.** Three-module support for the NPBM branch-control model. **a,** NPBM founder maps colored by BM-patterning, BL-selector and BL-reinforcement module scores. **b,** Founder module bias comparing $GLU^{BM}$-high and $GABA^{BL}$-high NPBM founders. $GLU^{BM}$-high founders start with higher BM-patterning signal, while $GABA^{BL}$-high founders show higher BL-selector signal. **c,** BL route module dynamics showing a shift from BL-selector signal toward BL-reinforcement signal over trajectory pseudo-time. **d,** Early and late BL local-control matrices summarizing inferred module-to-module influences. The early BL context shows strong support from BL reinforcement to BL selector and suppression of BM patterning, while the late BL context shows reinforcement self-support and continued suppression of the competing BM-patterning arm.

#### Supplementary Figure S13. Gene-program validation and timing of NPBM branch commitment

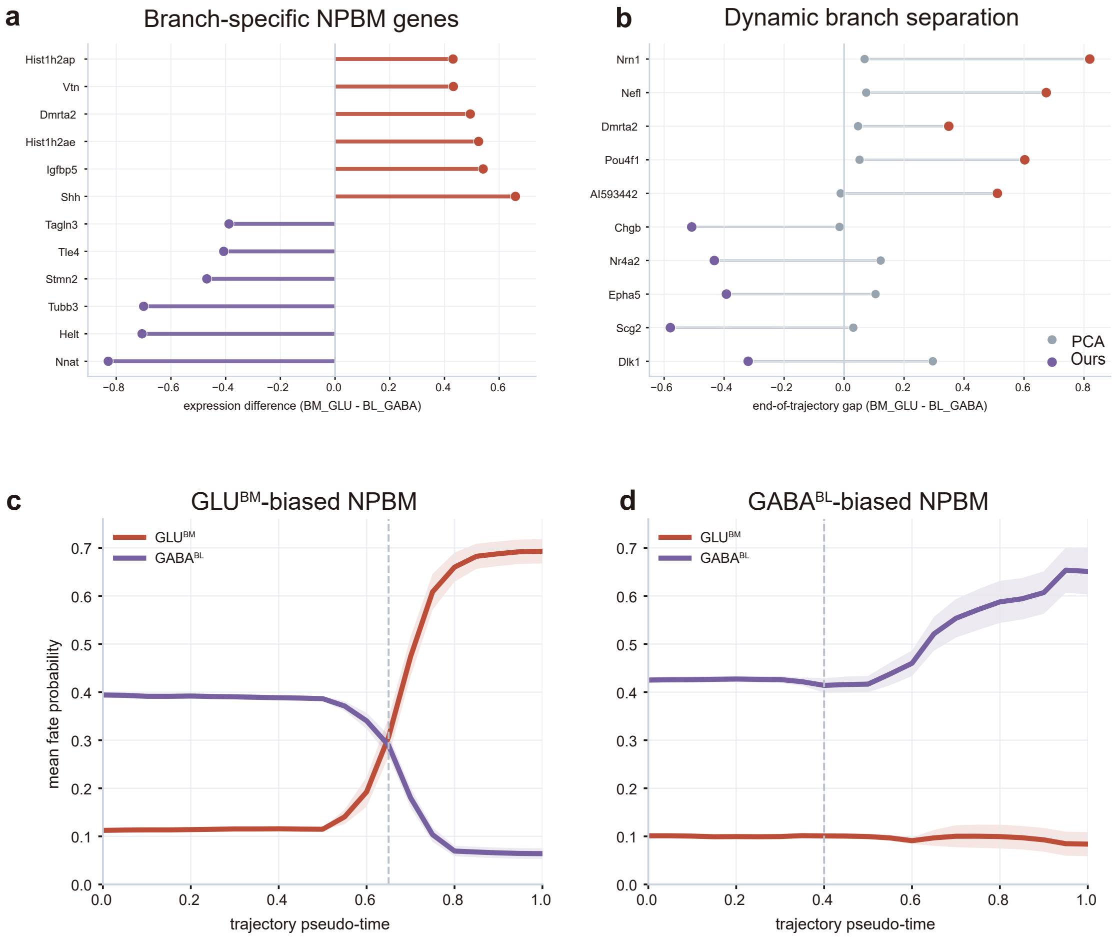


**Supplementary Figure S13.** Gene-program validation and timing of NPBM branch commitment. **a,** Branch-specific NPBM genes ranked by $\mathrm{GLU}^{\mathrm{BM}}$ minus $\mathrm{GABA}^{\mathrm{BL}}$ expression difference. **b,** Dynamic branch separation for representative genes at the end of inferred trajectories, comparing PhyloFM with a PCA baseline. **c,d,** Mean fate-probability trajectories for $\mathrm{GLU}^{\mathrm{BM}}$-biased and $\mathrm{GABA}^{\mathrm{BL}}$-biased NPBM founders. Dashed lines mark the inferred commitment window, where branch probabilities begin to separate sharply along the continuous rollout.

### Supplementary Figure S14. Founder bias and timing of the zebrafish fibroblast branch

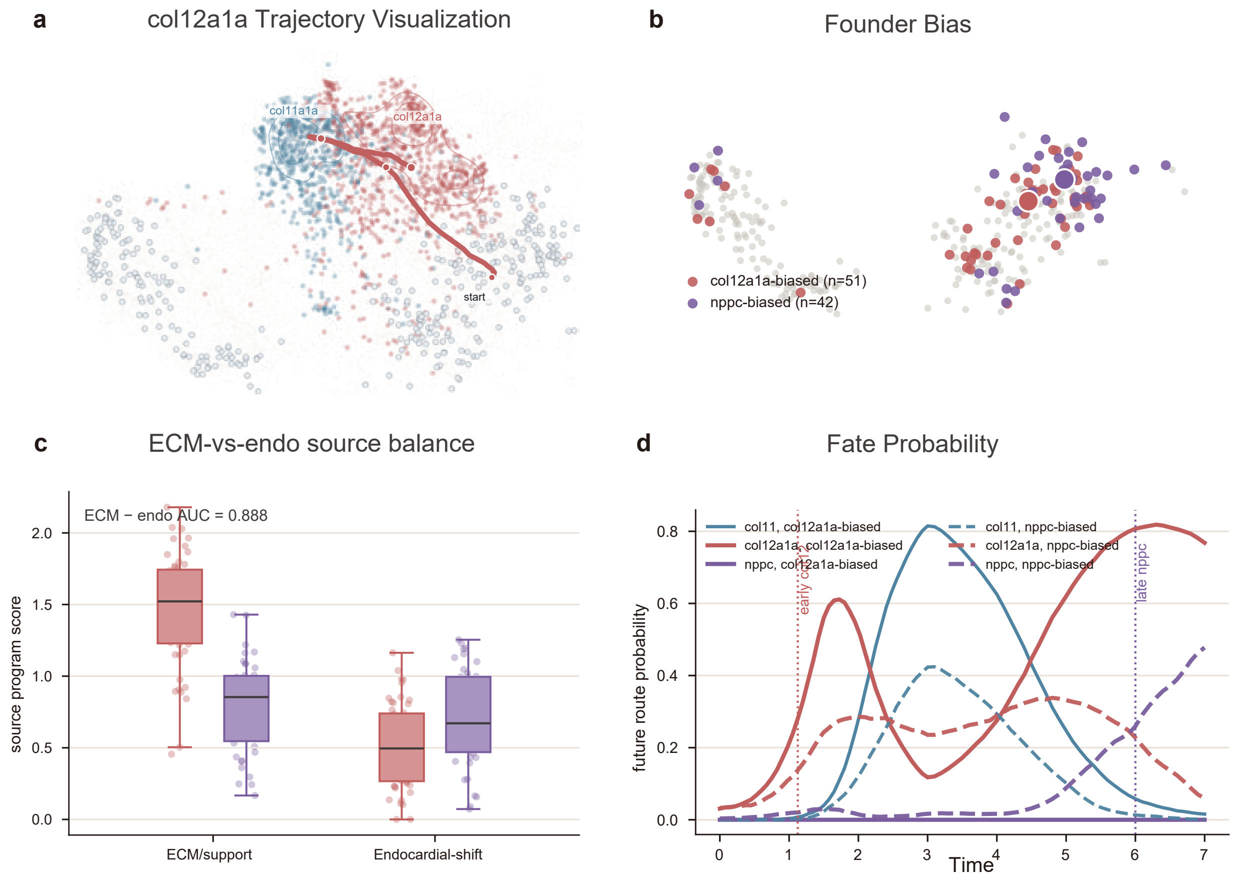


**Supplementary Figure S14.** Founder bias and timing of the zebrafish fibroblast branch. **a,** Visualization of the *col12a1a* trajectory through the fibroblast remodeling manifold. **b,** Founder-bias map for 0 dpi fibroblast sources, highlighting *col12a1a*-biased and *nppc*-biased founders. **c,** ECM-versus-endocardial source-program balance in *col12a1a*-biased and *nppc*-biased founders. The ECM minus endocardial-shift axis separates the two founder groups with AUC 0.888. **d,** Future route probabilities over model time for *col12a1a*-biased and *nppc*-biased founders. The *nppc* branch appears as a delayed diversion after an earlier *col12*-like phase.

### Supplementary Figure S15. Gene-program timing along zebrafish fibroblast routes

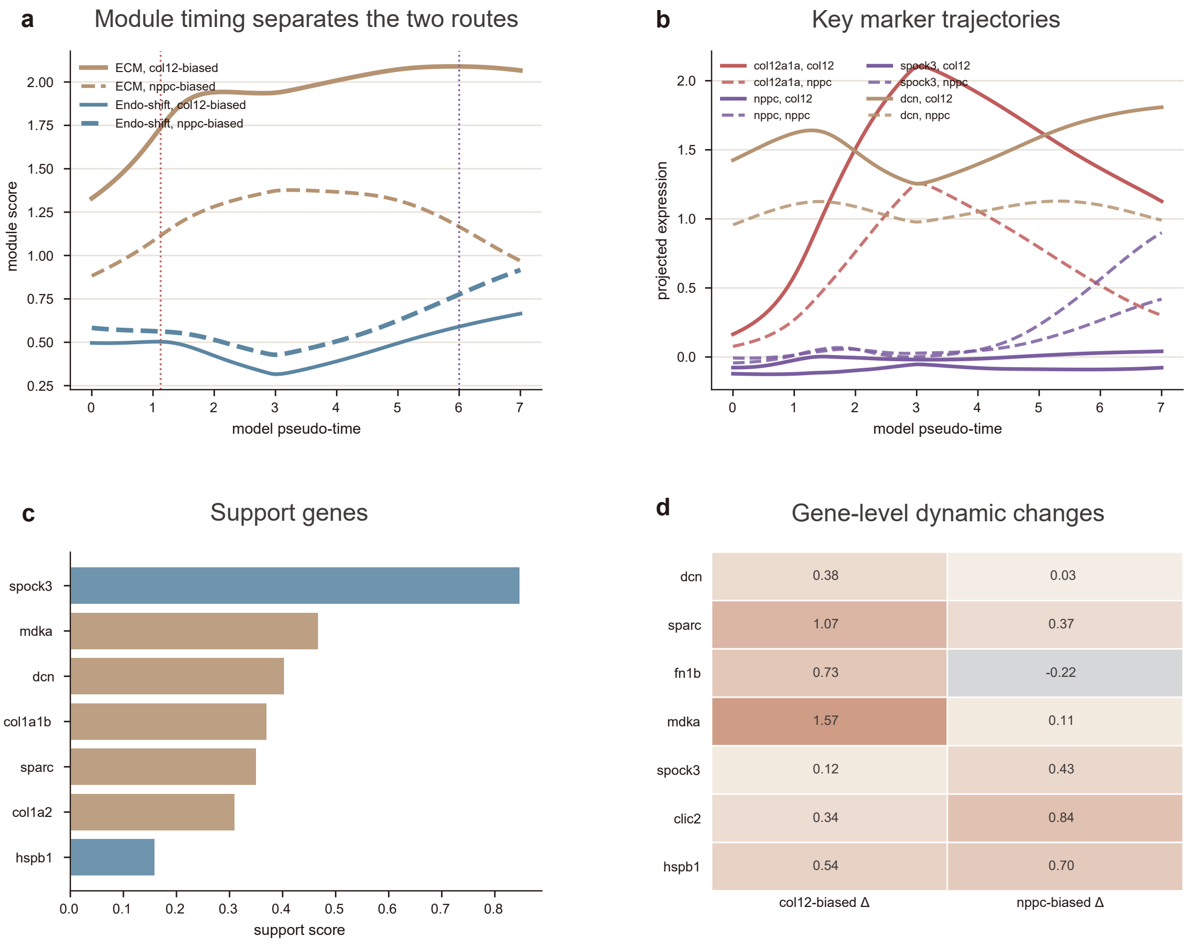


**Supplementary Figure S15.** Gene-program timing along zebrafish fibroblast routes. **a,** ECM and endocardial-shift module scores over model time in *col12a1a*-biased and *nppc*-biased trajectories. **b,** Marker-gene trajectories for representative ECM, *col12* and *nppc*-associated genes. **c,** Support genes ranked by branch-support score. **d,** Gene-level dynamic changes in *col12a1a*-biased and *nppc*-biased trajectories.

### Supplementary Figure S16. Velocity-GRN support for late *nppc* self-reinforcement

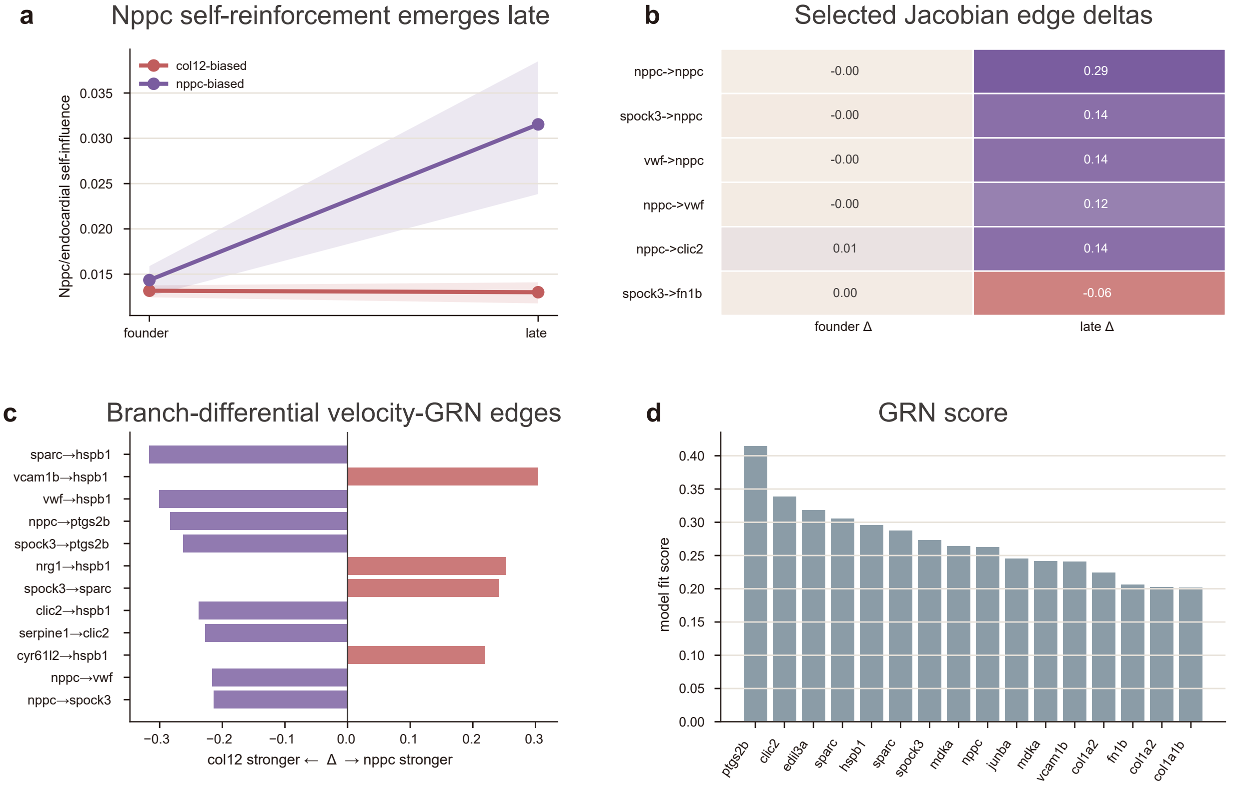


**Supplementary Figure S16.** Velocity-GRN support for late *nppc* self-reinforcement. **a,** *Nppc*/endocardial self-influence from founder to late trajectory states in *col12a1a*-biased and *nppc*-biased trajectories. **b,** Selected Jacobian edge deltas comparing founder and late branch states. **c,** Branch-differential velocity-GRN edges, with positive values stronger in the *nppc* branch and negative values stronger in the *col12a1a* branch. **d,** GRN score ranking for genes in the branch-specific velocity network.

#### Supplementary Figure S17. Secondary plasticity, inflammatory microstate and growth-ECM support analyses

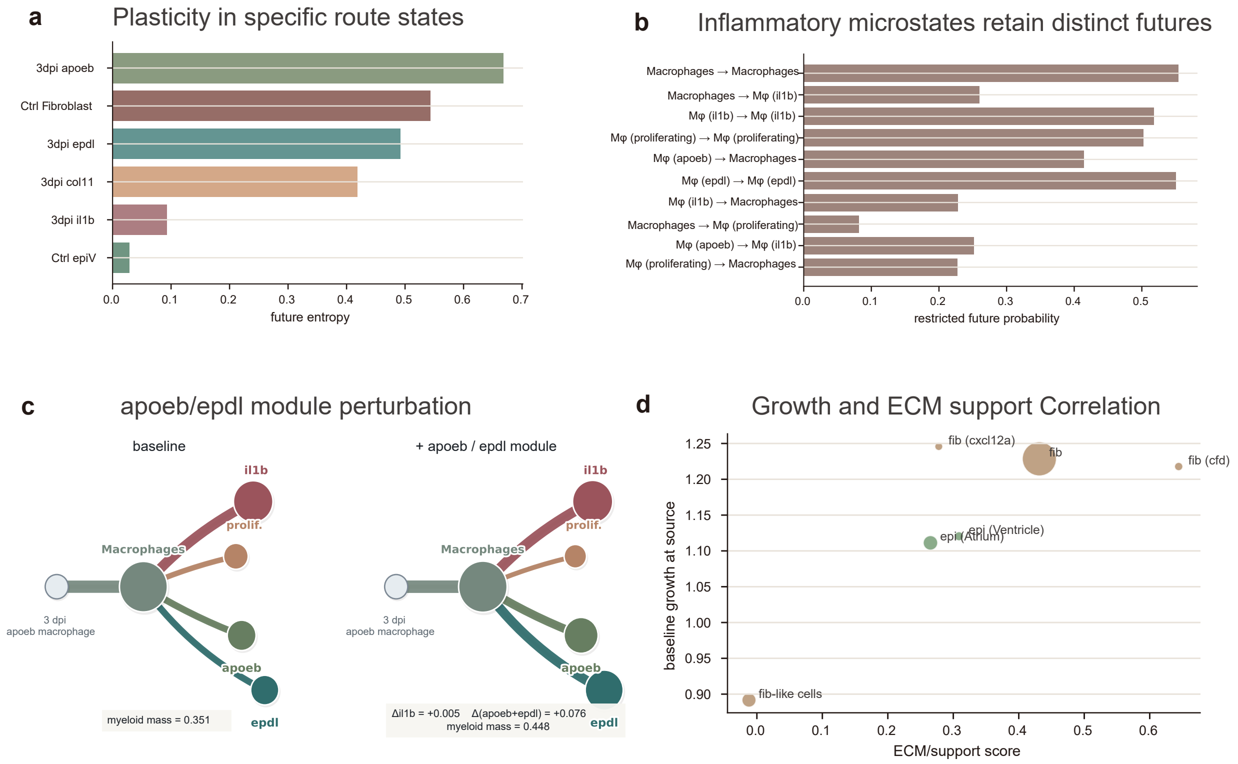


**Supplementary Figure S17.** Secondary plasticity, inflammatory microstate and growth-ECM support analyses. **a,** Future entropy across selected route states. **b,** Restricted future probabilities for inflammatory microstate transitions. **c,** Directional *apoeb*/*epdl* module perturbation support for inflammatory processing routes. **d,** Association between baseline growth at source and ECM-support score across fibroblast and epicardial support states.

### Supplementary Figure S18. Tumour-state transitions and module perturbation support in eTracer

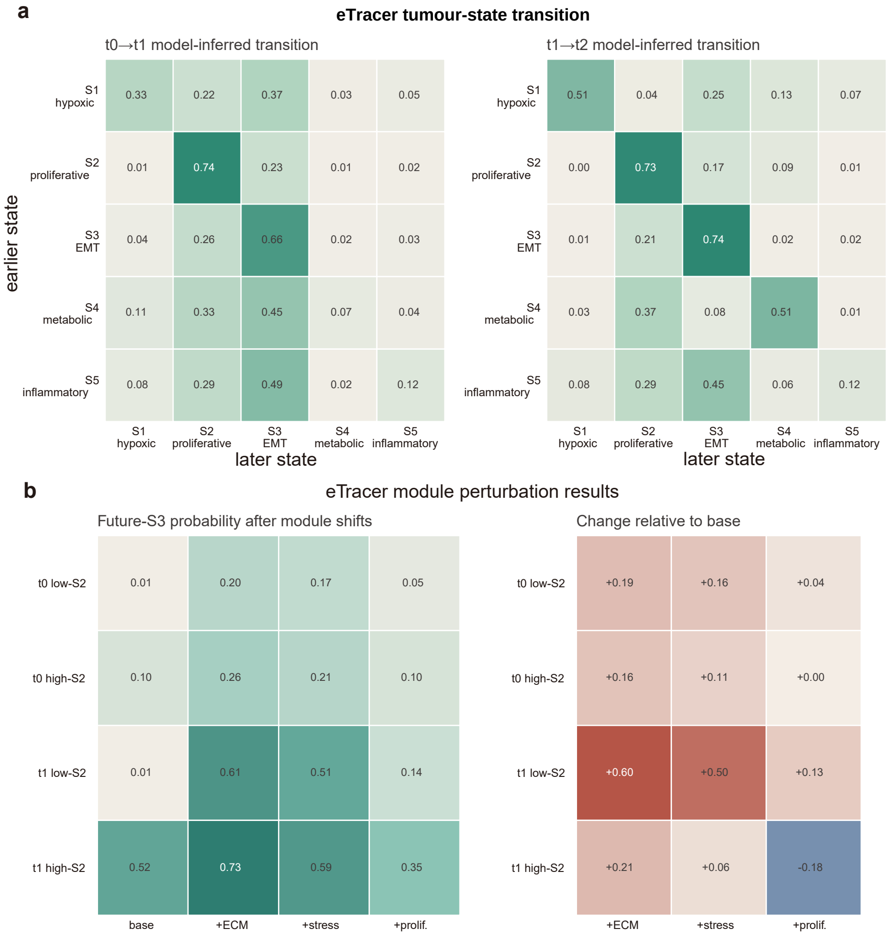


**Supplementary Figure S18.** Tumour-state transitions and module perturbation support in eTracer. **a,** Model-inferred tumour-state transition matrices for T0 to T1 and T1 to T2. Rows are earlier tumour states and columns are later tumour states. S2 remains self-retaining while continuing to transfer mass to S3, and S3 shows strong self-retention. **b,** Future-S3 probability after module shifts in low- and high-future S2 subsets. ECM and stress perturbations produce the strongest increases in future S3, especially in T1 low-S2 cells.

### Supplementary Figure S19. Continuous tumour-state composition across eTracer progression

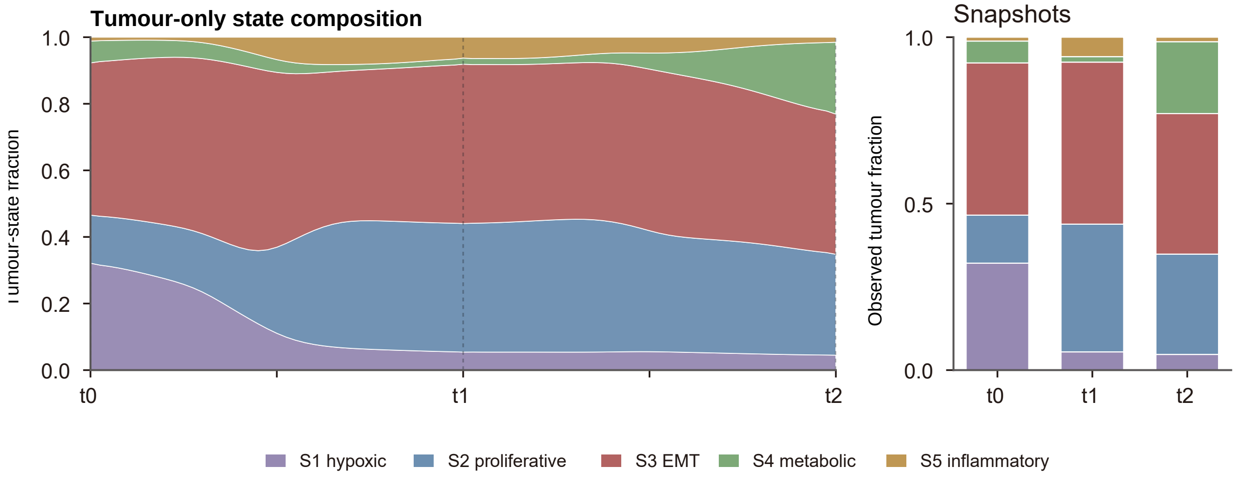


**Supplementary Figure S19.** Continuous tumour-state composition across eTracer progression. Left, model-inferred continuous tumour-state fractions from T0 to T2. Right, observed tumour-state fractions at the sampled snapshots. The continuous summary shows a decrease in S1 hypoxic fraction, persistent S2 proliferative mass and expansion of S3 EMT and later S4 metabolic fractions.

#### Supplementary Figure S20. Functional enrichment of future-S3-associated genes

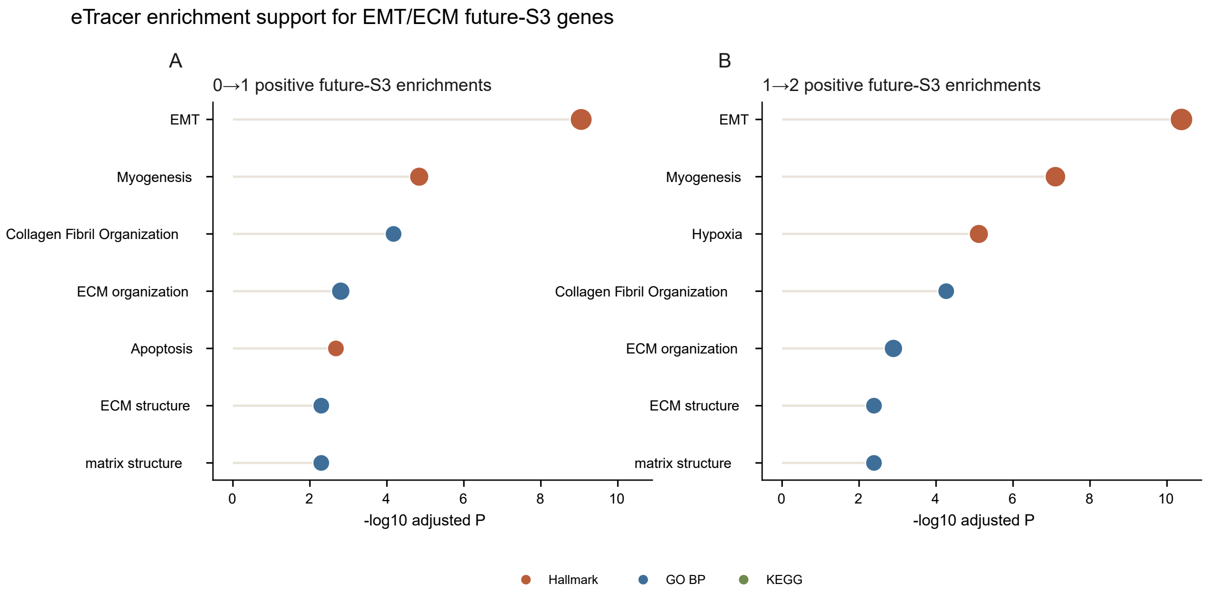


**Supplementary Figure S20.** Functional enrichment of future-S3-associated genes. **a,b,** Positive future-S3 gene enrichments for the 0 to 1 and 1 to 2 temporal intervals. Hallmark EMT, collagen fibril organization, extracellular matrix organization, ECM structure and matrix-structure terms are enriched across both intervals, supporting an EMT and matrix-remodeling program associated with future S3 competence.

[4] Marco Cuturi. Sinkhorn distances: Lightspeed computation of optimal transport. In *Advances in Neural Information Processing Systems*, volume 26, 2013.

[10] Zhenyi Zhang, Tiejun Li, and Peijie Zhou. Learning stochastic dynamics from snapshots through regularized unbalanced optimal transport. In *The Thirteenth International Conference on Learning Representations*, 2025.

[11] Yuhao Sun, Zhenyi Zhang, Zihan Wang, Tiejun Li, and Peijie Zhou. Variational regularized unbalanced optimal transport: Single network, least action. *Advances in Neural Information Processing Systems*, 2025.

[12] Dongyi Wang, Yuanwei Jiang, Zhenyi Zhang, Xiang Gu, Peijie Zhou, and Jian Sun. Joint velocity-growth flow matching for single-cell dynamics modeling. *Advances in Neural Information Processing Systems*, 2025.

[13] Zhenyi Zhang, Zihan Wang, Yuhao Sun, Tiejun Li, and Peijie Zhou. Modeling cell dynamics and interactions with unbalanced mean field Schrödinger Bridge. *Advances in Neural Information Processing Systems*, 2025.

[14] Yaron Lipman, Ricky T. Q. Chen, Heli Ben-Hamu, Maximilian Nickel, and Matthew Le. Flow matching for generative modeling. In *The Eleventh International Conference on Learning Representations*, 2023.

[15] Alexander Tong, Kilian Fatras, Nikolay Malkin, Guillaume Huguet, Yanlei Zhang, Jarrid Rector-Brooks, Guy Wolf, and Yoshua Bengio. Improving and generalizing flow-based generative models with minibatch optimal transport. *Transactions on Machine Learning Research*, 2024.

[23] Peter N. Inglis, Guangshuo Ou, Michel R. Leroux, and Jonathan M. Scholey. The sensory cilia of Caenorhabditis elegans. In *WormBook*. The C. elegans Research Community, 2007.

[53] Mehrshad Sadria, Allen Zhang, and Gary D. Bader. Deep Lineage: Single-cell lineage tracing and fate inference using deep learning. *bioRxiv*, page 2024.04.25.591126, 2024.

[54] Sergey Isaev, Alek G. Erickson, Igor Adameyko, and Peter V. Kharchenko. Clonal embeddings allow exploratory analysis of lineage-resolved single-cell data. *bioRxiv*, page 2026.04.30.720820, 2026.

[55] John Isaac Murray et al. Multidimensional regulation of gene expression in the C. elegans embryo. *Genome Research*, 22(7):1282–1294, 2012.
